# The temporal progression of immune remodeling during metastasis

**DOI:** 10.1101/2023.05.04.539153

**Authors:** Christopher S. McGinnis, Zhuang Miao, Nathan E. Reticker-Flynn, Juliane Winker, Ansuman T. Satpathy

## Abstract

Tumor metastasis requires systemic remodeling of distant organ microenvironments which impacts immune cell phenotypes, population structure, and intercellular communication networks. However, our understanding of immune phenotypic dynamics in the metastatic niche remains incomplete. Here, we longitudinally assayed lung immune cell gene expression profiles in mice bearing PyMT-driven metastatic breast tumors from the onset of primary tumorigenesis, through formation of the pre-metastatic niche, to the final stages of metastatic outgrowth. Computational analysis of these data revealed an ordered series of immunological changes that correspond to metastatic progression. Specifically, we uncovered a TLR-NFκB myeloid inflammatory program which correlates with pre-metastatic niche formation and mirrors described signatures of CD14+ ‘activated’ MDSCs in the primary tumor. Moreover, we observed that cytotoxic NK cell proportions increased over time which illustrates how the PyMT lung metastatic niche is both inflammatory and immunosuppressive. Finally, we predicted metastasis-associated immune intercellular signaling interactions involving *Igf1* and *Ccl6* which may organize the metastatic niche. In summary, this work identifies novel immunological signatures of metastasis and discovers new details about established mechanisms that drive metastatic progression.

**Graphical abstract:** 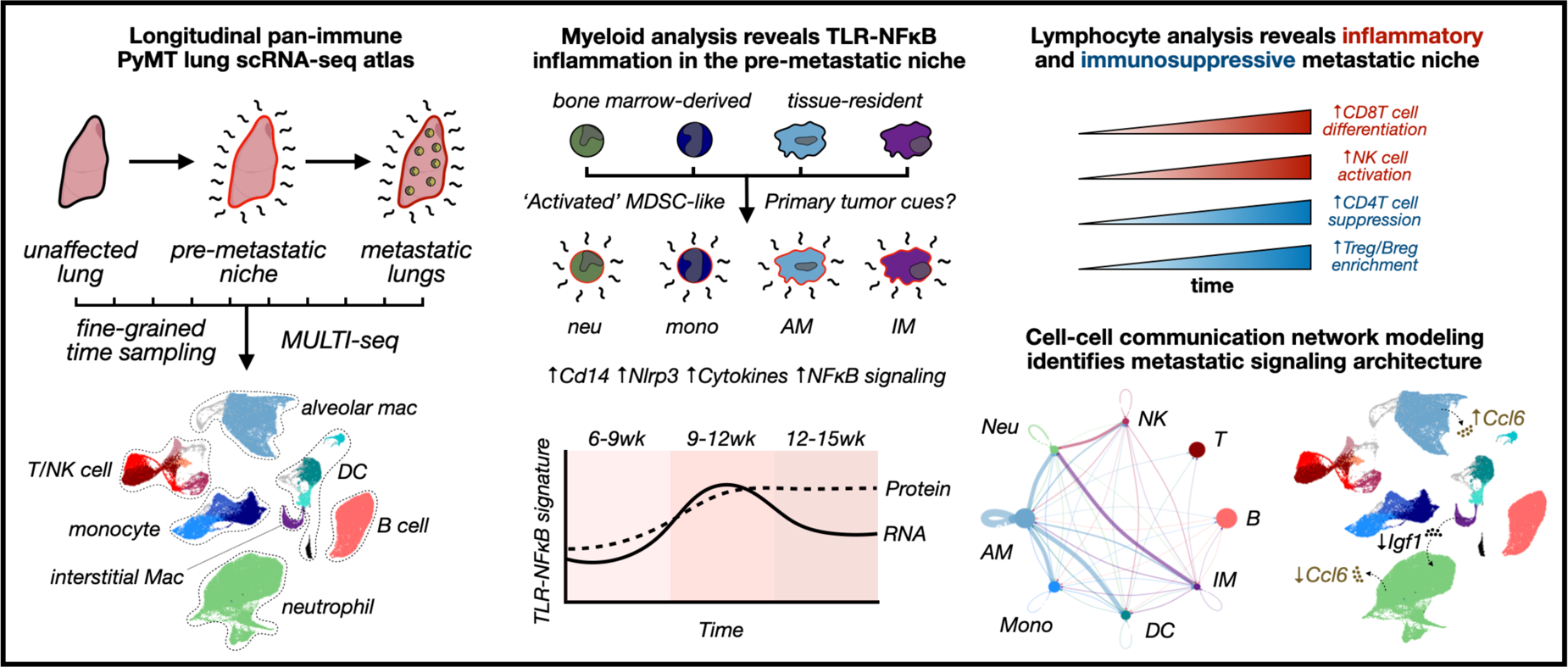

**In brief:** McGinnis et al. report a longitudinal scRNA-seq atlas of lung immune cells in mice bearing PyMT-driven metastatic breast tumors and identify immune cell transcriptional states, shifts in population structure, and rewiring of cell-cell signaling networks which correlate with metastatic progression.

**Highlights:**

- Longitudinal scRNA-seq reveals distinct stages of immune remodeling before, during, and after metastatic colonization in the lungs of PyMT mice.
- TLR-NFκB inflammation correlates with pre-metastatic niche formation and involves both tissue-resident and bone marrow-derived myeloid cell populations.
- Inflammatory lung myeloid cells mirror ‘activated’ primary tumor MDSCs, suggesting that primary tumor-derived cues induce *Cd14* expression and TLR-NFκB inflammation in the lung.
- Lymphocytes contribute to the inflammatory and immunosuppressive lung metastatic microenvironment, highlighted by enrichment of cytotoxic NK cells in the lung over time.
- Cell-cell signaling network modeling predicts cell type-specific *Ccl6* regulation and IGF1-IGF1R signaling between neutrophils and interstitial macrophages.

## INTRODUCTION

Cancer metastasis is the cause of death for 50-90% of patients with solid tumors^1^ in part because clinically-approved therapies which specifically target metastasis do not exist. Prior work has demonstrated that systemic tumor-mediated immune remodeling is necessary for metastatic progression.^2–7^ As a result, specifically targeting metastatic disease by reversing or reorienting immune dysregulation in the metastatic niche of distant organ sites represents an appealing yet relatively unexplored strategy. However, since most preclinical studies of tumor-immune interactions have relied upon cross-sectional analyses of immune cell phenotypes in primary tumors and unaffected tissues, our understanding of how immune cell populations are dynamically remodeled in the metastatic niche remains incomplete.

Pro-metastatic immune remodeling involves diverse changes in immune cell state (e.g., induction of immunosuppressive myeloid phenotypes^8, 9^), population structure (e.g., ‘left-shift’ bias towards myelopoiesis^9^), trafficking patterns (e.g., selective recruitment of cells to the metastatic microenvironment^11^), and intercellular communication networks (e.g., increased expression of myeloid chemoattractants such as *Ccl2*^6^). High-throughput single-cell genomics analysis can simultaneously assay these critical immunological signatures, as has been described for late-stage metastatic disease.^3, 4^ However, since the metastatic niche is dynamically reorganized during disease progression,^12, 13^ longitudinal single-cell genomics analyses of immune remodeling in distant organ sites during metastasis has the potential to both refine our understanding of metastatic immune remodeling and identify candidate pathways for anti-metastatic immunotherapy development.

To this end, we constructed a longitudinal single-cell RNA-sequencing (scRNA-seq) cell atlas of lung immune cells in the polyomavirus middle T antigen (PyMT) mouse model of spontaneous metastatic breast cancer. Unlike prior scRNA-seq analyses of metastatic niche immune cells in experimental metastasis^14, 15^ and patient-derived xenograft^16^ models, our atlas systematically chronicles how the fully-intact metastatic niche immune compartment responds during primary tumorigenesis, through pre-metastatic niche formation, to the final stages of metastatic outgrowth in the lung. In-depth computational analysis of these data yielded three key insights into tumor-mediated immune reprogramming in PyMT mice.

First, analyses of lung myeloid cells revealed that bone marrow (BM) derived and tissue-resident cell types in the pre-metastatic niche engage in toll-like receptor (TLR) inflammatory signaling through the NFκB pathway. While ‘sterile’ TLR-NFκB inflammation plays a known role in metastatic progression,^17–21^ our analyses identify cell types that engage in TLR-NFκB inflammation which were not previously known and suggest that primary tumor-derived cues act at a distance to reprogram lung myeloid cells to participate in this process. Second, analyses of lung lymphoid cells identified a previously unobserved increase in cytotoxic natural killer (NK) cell proportions over time which highlights how the PyMT metastatic microenvironment is simultaneously inflammatory and immunosuppressive. Third, cell-cell signaling network modeling predicted metastasis-associated interactions such as IGF1-IGF1R and CCL6-CCR1/CCR2 signaling which may organize the metastatic niche and represent candidate pathways for anti-metastatic immunotherapy development. Considered collectively, our longitudinal scRNA-seq atlas of PyMT lung immune cells revealed novel signatures of metastatic immune remodeling and represents a critical resource for development of anti-metastatic immunotherapies.

## RESULTS

### A longitudinal scRNA-seq cell atlas of PyMT mouse lungs reveals immune population structure dynamics in the metastatic microenvironment

PyMT mice are a well-established model of metastatic breast cancer that is caused by mammary-specific expression of the polyomavirus middle T antigen transgene driven by the murine mammary tumor virus long terminal repeat promoter.^22^ PyMT+ female mice develop mammary tumors by 6-8 weeks of age with 100% tumor penetrance and 80-90% of tumor-bearing mice develop lung metastases by 14-16 weeks.^22, 23^ PyMT tumors are transcriptionally similar to the luminal B subtype of human breast cancer,^24^ and PyMT primary tumor development mimics many features of human disease.^25^ PyMT is an ideal model for temporal analyses of immune remodeling during metastatic progression because of the well-studied disease progression dynamics and high tumor penetrance, combined with recapitulation of the entire metastatic cascade and presence of an intact immune system in the autochthonous model.

To longitudinally profile immune cell transcriptional changes in the metastatic niche, we used the single-cell genomics sample multiplexing technology we previously developed, MULTI-seq,^16^ to construct a scRNA-seq atlas of CD45+ immune cells isolated from PyMT+ female mice weekly between 6-15 weeks of age and 9-week-old wild-type (WT) controls (**Fig. 1A**; Experimental Methods). By tagging cells with sample-specific MULTI-seq barcodes prior to pooled scRNA-seq, our workflow enabled us to ‘super-load’ droplet microfluidic lanes to increase cell-throughput, computationally identify and remove doublets, and avoid potentially-confounding batch effects while analyzing timepoints spanning the onset of primary tumorigenesis to the final stages of metastatic outgrowth. Following MULTI-seq sample demultiplexing, doublet removal, and quality-control filtering (Computational Methods), we proceeded with the analysis of 104,314 high-quality immune cells. All week 6-14 timepoints and controls were represented by at least 2 biological replicates and each sample included an average of 2,749 cells (**Fig. S1A**).

**Figure 1:**
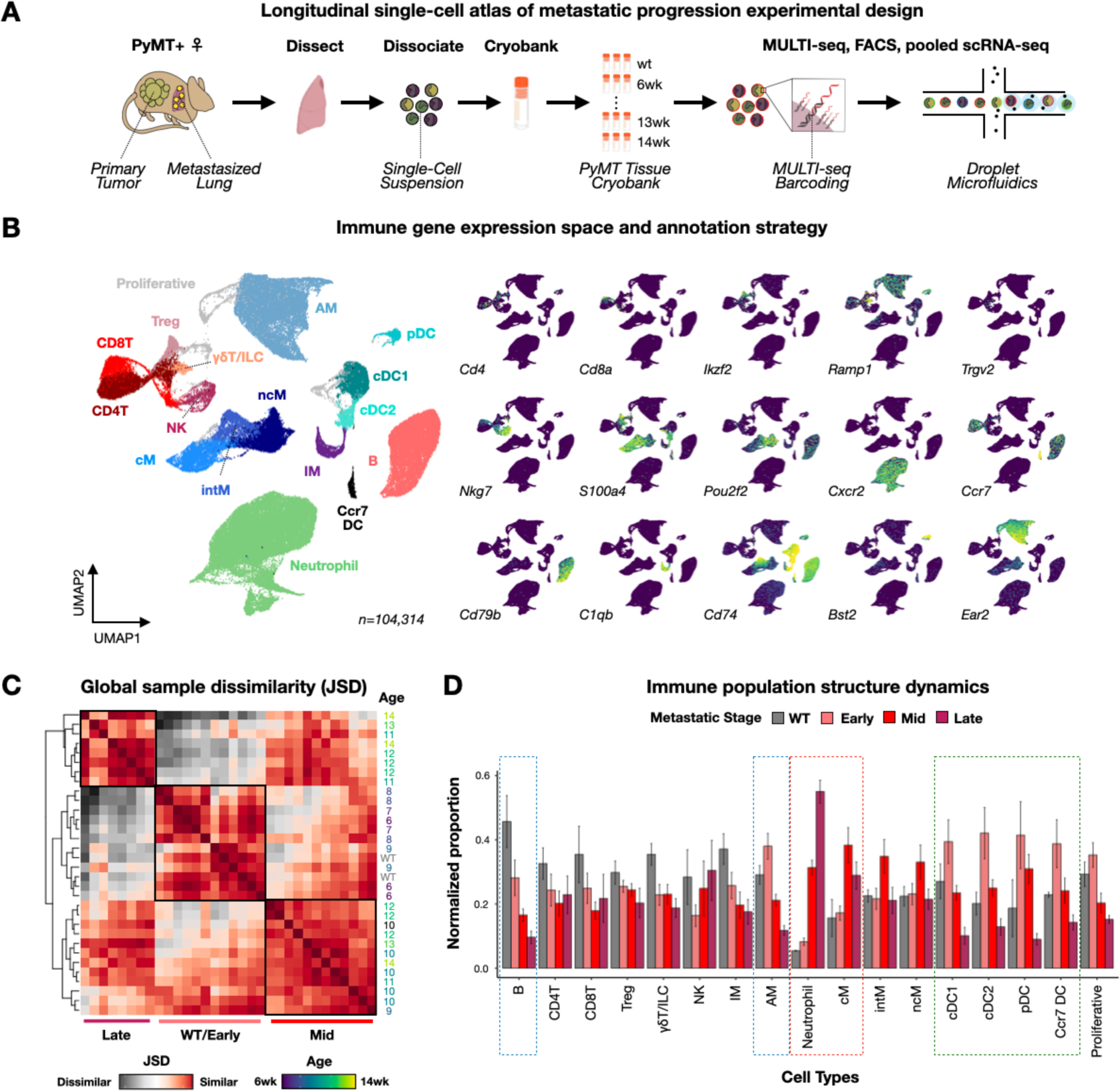
Longitudinal scRNA-seq cell atlas of PyMT mouse lung immune cells captures dynamics of the metastatic microenvironment. (A) Schematic of experimental approach. (B) Visualization of immune gene expression space with feature plots of annotation markers for major immune cell types. (C) JSD heatmap demonstrating (dis-)similarity between PyMT lung samples. Hierarchical clustering-defined clades representing WT/early-stage metastasis (light red), mid-stage (red), and late-stage (maroon) highlighted. Week 15 PyMT mouse sample excluded. (D) Bar charts describing shifts in immune cell type proportions during metastatic progression. Progressively-enriched, progressively-depleted, and transiently-enriched cell types highlighted with red, blue, and green boxes, respectively. Data represented as per cell type normalized mean ± s.e.m.

Cell type annotation using unsupervised clustering and differential gene expression analysis identified all expected major immune cell types including four T cell lineages: CD4 and CD8 T cells, regulatory T cells (Treg), and γδ T cells (γδT); three monocyte subsets: classical (cM), non-classical (ncM), and intermediate (intM); two types of tissue-resident macrophages: interstitial (IM) and alveolar (AM); four types of dendritic cells (DC): *Ccr7*+ DC, type 1 conventional DC (cDC1), type 2 conventional DC (cDC2), and plasmacytoid DC (pDC); as well as neutrophils, NK, B, proliferative, and innate lymphoid cells (ILC; **Fig. 1B, Fig. S1B, Table S1**). We next sought to benchmark our data against known changes in immune population structure during metastatic progression. Due to natural variability in the timing of disease progression, we opted against modeling experimental timepoints continuously. Instead, we used the dissimilarity metric Jensen-Shannon Divergence (JSD)^26, 27^ to computationally group samples into three metastatic stages in an unbiased manner: (1) ‘early-stage’ lungs from WT and 6-9 week-old mice, (2) ‘mid-stage’ lungs primarily from 9-12 week-old mice, and (3) ‘late-stage’ lungs primarily from 12-14 week-old mice (**Fig. 1C**). Since 9-12 week old PyMT+ mice harbor large primary breast tumors which have not yet metastasized to the lung,^22, 23^ we refer to mid-stage lungs as the pre-metastatic niche in this study.

Comparing normalized cell type proportions between JSD-defined stages demonstrated that the frequencies of neutrophils and cMs increased during metastatic progression while AM frequencies decreased (**Fig. 1D**), matching prior reports in multiple model systems^28–30^ and, thus, successfully benchmarking our JSD-defined metastatic stage groupings. Notably, we also observed metastasis-associated reductions in B cell frequencies, as well as transient enrichment of DC proportions in early-stage lungs which would have been missed without the fine-grained longitudinal dimension of our scRNA-seq atlas. These insights illustrate the utility of our approach in measuring the dynamics of immune remodeling during metastasis.

### Tissue-resident macrophages enact wound healing and TLR-NFκB inflammation responses in the pre-metastatic niche

We next focused on analyzing lung myeloid cells since metastasis is known to require profound tumor-mediated remodeling of myeloid cell lineages.^28, 31^ Moreover, experimental manipulation of different myeloid cell types can influence metastatic progression via diverse mechanisms,^32–37^ which positions these cells as potential targets for anti-metastatic immunotherapy development. We identified two main groups of myeloid cells in our data: (1) BM-derived monocytes, neutrophils, and DCs which migrate into the lung and (2) tissue-resident macrophages such as IMs and AMs.^38^ While the roles of most BM-derived myeloid cells in metastasis have been studied extensively,^12, 31^ how IMs and AMs contribute to metastatic progression is relatively under-studied.

In IMs, we distinguished three main subtypes (**Fig. 2A**, left; **Fig. S2A; Table S2**) including *Mrc1*+ IMs and *Cd74*+ antigen-presenting IMs which were previously seen in tumor-naïve mice,^39–41^ as well as *Crip1*+ IMs which were not observed in tumor-naïve or lung-metastatic 4T1 tumor-bearing mice.^14^ *Crip1*+ IM proportions were modestly increased in mid-stage pre-metastatic niche lungs (**Fig. 2A**, right) and shared many transcriptional features with *Crip1*^hi^ *Cav1*+ macrophages observed in mouse models of knee damage^42^ (**Fig. 2B**), suggesting that *Crip1*+ IMs may contribute to tissue damage responses in PyMT mouse lungs.

**Figure 2.**
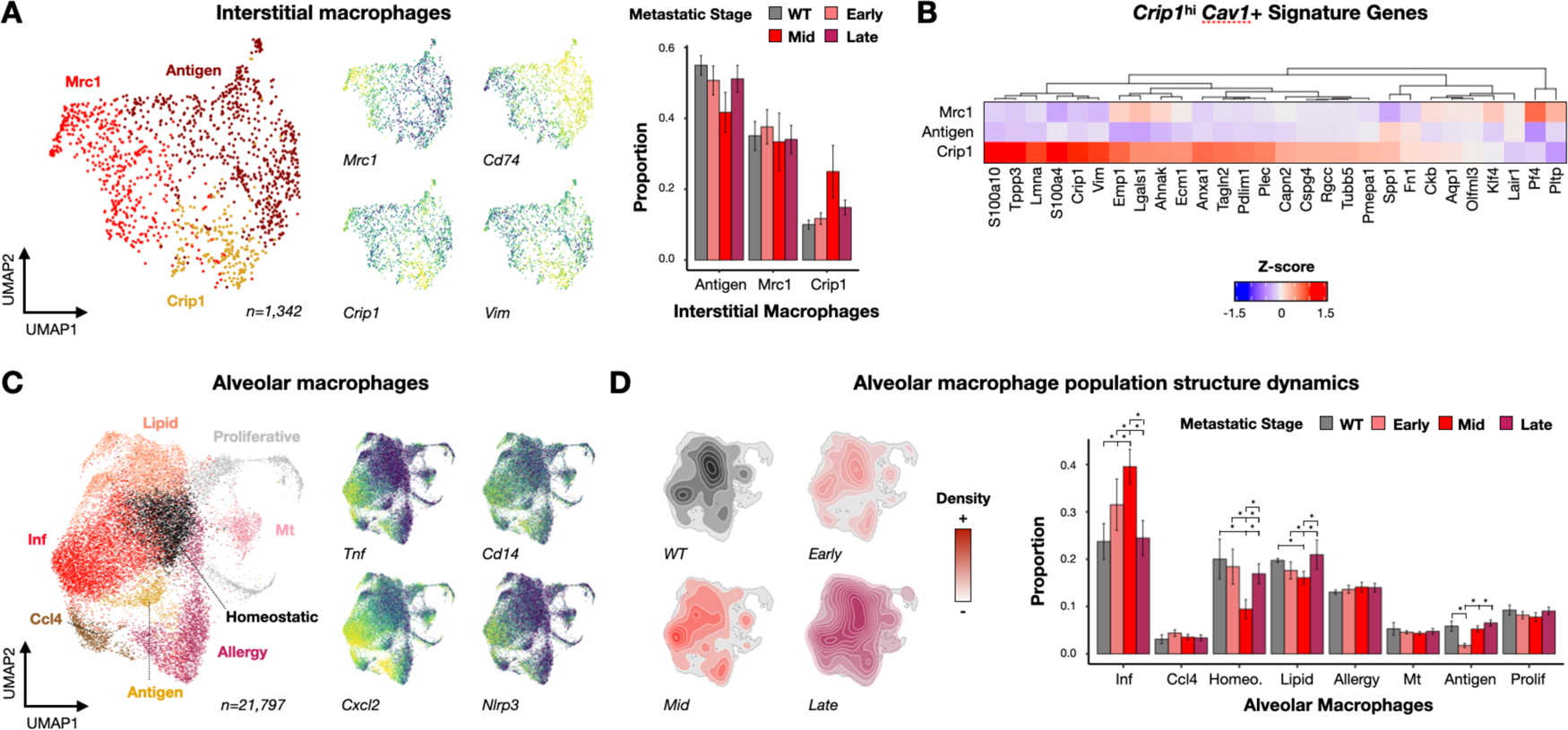
AM inflammation and IM wound healing transcriptional signatures are linked to pre-metastatic niche formation. (A) Visualization of IM gene expression space colored by subtype with feature plots of subtype markers and bar charts describing shifts in IM proportions during metastatic progression. Proportional data represented as mean ± s.e.m. * corresponds to p < 0.01 in two-proportion z-tests. (B) Z-score heatmap of *Crip1*^hi^ *Cav1*+ signature genes in IM subtypes. (C) Visualization of AM gene expression space colored by subtype with feature plots for *Cd14*+ inflammatory AM cell markers. (D) Visualization of metastatic stage densities in AM gene expression space (left) with bar charts describing shifts in AM proportions over time. Proportional data represented as mean ± s.e.m. * corresponds to p < 0.01 in two-proportion z-tests.

In AMs, we identified eight total subtypes, five of which reflect known AM functions^43^ such as antigen presentation, allergic response, surfactant homeostasis, *Ccl4*+ inflammation, and proliferation (**Fig. 2C**, left; **Fig. S2B; Table S2**). We also identified two disease-associated subtypes including ‘lipid-associated’ AMs that were previously described in 4T1 tumor-bearing mice^14^ and metallothionein (Mt) expressing AMs which were also detected in a mouse model of chronic obstructive pulmonary disease.^44^ Finally, we detected a novel subtype of inflammatory AMs expressing the cytokines *Tnf* and *Cxcl2*, the inflammasome component *Nlrp3*, and the TLR co-receptor *Cd14* (**Fig. 2C**, right).^45^ Compared to other AM subtypes, *Cd14*+ inflammatory AMs were enriched in pre-metastatic niche lungs while surfactant homeostasis AM frequencies decreased (**Fig. 2D**), suggesting that pre-metastatic niche formation in PyMT tumor-bearing mice involves the induction of an inflammatory response in these cells.

Lung immune subtypes that correlate with pre-metastatic niche formation represent attractive targets for the development of anti-metastatic immunotherapies. Thus, we used non-negative matrix factorization (NMF)^46^ to identify a detailed transcriptional signature for *Cd14*+ inflammatory AMs. NMF revealed a single component (NMF25) that exhibited enrichment not only in *Cd14*+ inflammatory AMs (**Fig. 3A**, left), but also subpopulations of each AM subtype, as was highlighted by sub-clustering analysis after removing *Cd14*+ inflammatory and surfactant homeostasis AMs (**Fig. 3A**, middle and right). These sub-clustered inflammatory subsets also expressed *Cd14*, *Cxcl2*, and *Nlrp3* (**Fig. 3B**) and were enriched in pre-metastatic niche lungs (**Fig. 3C**). Gene set enrichment analysis (GSEA)^47^ comparing NMF25 signature genes (**Table S3**) to Hallmark gene sets^48^ revealed enrichment for many inflammatory pathways including TNF signaling via NFκB (**Fig. 3D**). Indeed, many genes up-regulated by inflammatory AMs include NFκB signaling components, which is consistent with active CD14-mediated TLR-NFκB signaling through the MyD88-dependent pathway^45, 49^ by inflammatory AMs.

**Figure 3.**
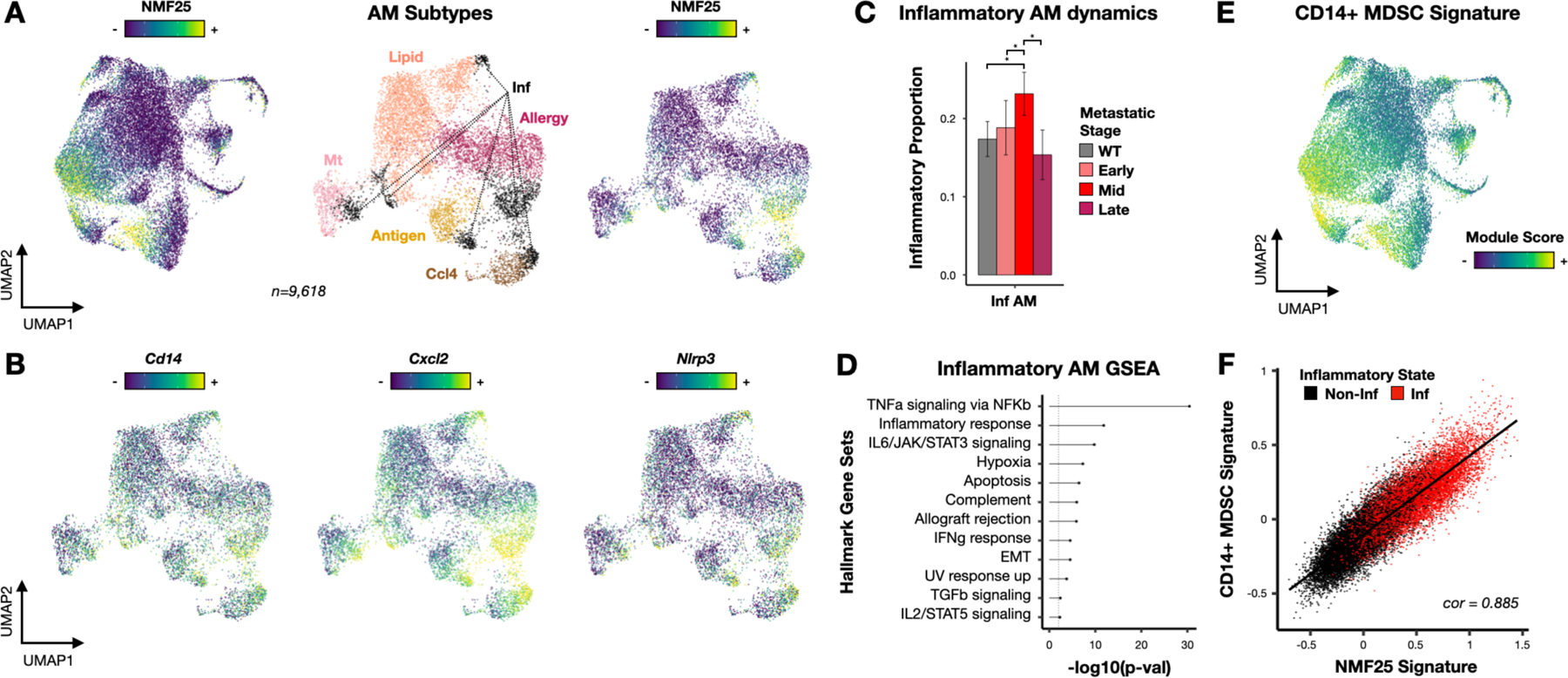
NMF and GSEA links *Cd14*+ inflammatory AM signature to TLR-NFκB inflammation and CD14+ ‘activated’ MDSCs. (A) Visualization of AM gene expression space before (left) and after sub-clustering (middle and right) colored by NMF25 score or subtype. (B) Visualization of sub-clustered AM gene expression space colored by TLR-NFκB inflammation signature gene expression. (C) Bar charts describing shifts in *Cd14*+ inflammatory AM proportions over time. * corresponds to p < 0.01 in two-proportion z-tests. (D) Lollipop plot depicting Hallmark gene sets that are significantly enriched amongst genes comprising the NMF25 transcriptional signature. (E) Visualization of AM gene expression space colored by CD14+ ‘activated’ MDSC module score. (F) Scatter plot illustrating correlation between CD14+ ‘activated’ MDSC and NMF25 module scores in AMs. Cells colored by inflammation state.

TLR-NFκB inflammatory signaling in lung epithelial cells^21^ and immunosuppressive BM-derived myeloid cells^17–20^ is known to promote lung metastasis. However, involvement of AMs in this process has not been described. Moreover, while primary tumors induce the expression of endogenous TLR ligands in distant organ sites,^17–20^ cues which drive expression of TLRs and the co-receptor CD14 in the metastatic niche are not known. Recent single-cell analyses of splenic and primary tumor neutrophils from multiple orthotopic transplant models have shown that myeloid derived suppressor cell (MDSC) ‘activation’ in the primary tumor is associated with increased *Cd14* expression and TLR-NFκB inflammation.^50^ Comparing gene expression module scores between NMF25 and the ‘activated’ MDSCs profiled in this study revealed significant alignment (**Fig. 3E; Fig. 3F**). This result suggests that the TLR-NFκB inflammatory program we observe in tissue-resident AMs may be induced by the same primary tumor-derived cues which locally ‘activate’ MDSCs.

### BM-derived myeloid cells contribute to pre-metastatic niche TLR-NFκB inflammation

We next turned our focus to the BM-derived myeloid lineages: monocytes, DCs, and neutrophils. Metastatic niche involvement of BM-derived myeloid cells has been studied extensively^12, 31^ and signatures of many known effects were validated in our scRNA-seq atlas. For instance, in monocytes we identified reciprocal shifts in ncM and cM proportions,^8, 51^ as well as monocytic MDSCs (M-MDSCs) that were exclusively linked to late-stage 15-week old PyMT mice (**Fig. 4A; Fig. S3A; Table S4**). Moreover, we observed that mature neutrophils enacted a degranulation-related transcriptional signature^52–55^ in mid- and late-stage lungs (**Fig. S4**), and that immature and transitional neutrophils were proportionally enriched relative to mature neutrophils over time (**Fig. 4B**, top; **Fig. S3B; Table S4**).^9, 10, 29^

**Figure 4.**
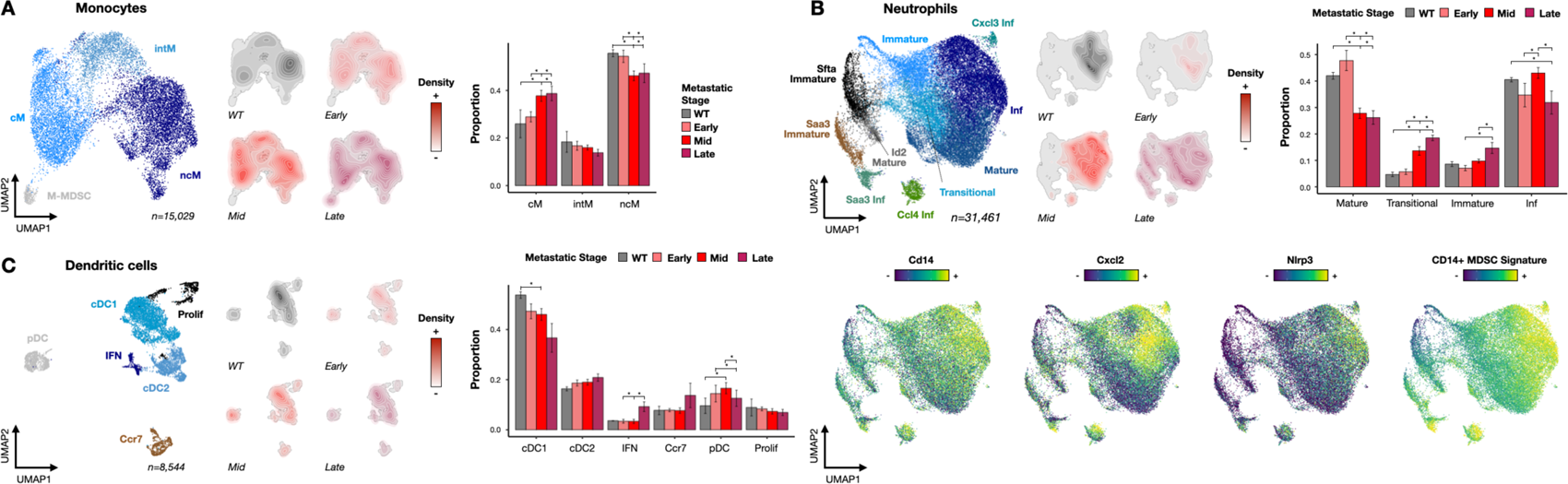
BM-derived myeloid subtype characterization uncovers metastasis-associated DC subtype frequency shifts and induction of TLR-NFκB inflammation in pre-metastatic niche neutrophils. (A) Visualization of monocyte gene expression space colored by subtype annotations (left) or metastatic stage densities (middle) with bar charts describing shifts in monocyte subtype proportions during metastatic progression (right). M-MDSCs were excluded from subtype proportion visualization for clarity due to exclusivity to late-stage samples. Proportional data represented as mean ± s.e.m. * corresponds to p < 0.01 in two-proportion z-tests. (B) Neutrophil analyses as presented in Fig. 4A (top) with feature plots of TLR-NFκB inflammation signature gene expression (bottom). Neutrophil subsets exclusively linked to late-stage samples were excluded from subtype proportion visualization for clarity. (C) DC analyses as presented in Fig. 4A.

Beyond these established immunological signatures, we also found metastasis-associated shifts in DC and neutrophil subtype distributions and phenotypes. For instance, we detected interferon-stimulated (IFN) and proliferative DCs which likely reflect responses to inflammation since DC proliferation in tissues is limited under steady-state conditions (**Fig. 4C; Fig. S3C; Table S4**).^56^ Moreover, we observed that cDC1 proportions were depleted over time (as described previously in metastasized PyMT lymph nodes^3^) while pDCs were enriched in pre-metastatic niche lungs (**Fig. 4C**, right) – two results which are consistent with the known biology of cDC1 promotion of anti-tumor immunity^31, 57^ and pDC recruitment of pro-metastatic cell types such as MDSCs and Tregs.^58, 59^ In neutrophils, we detected subsets expressing the TLR ligand *Saa3*^17^ or genes associated with neutrophil differentiation (*Id2*^60, 61^) and tissue damage (*Sfta2*^62^; **Fig. 4B**, top; **Fig. S3B**) which were exclusively linked to 15-week PyMT mice (**Fig. S5**), suggesting that neutrophils uniquely respond to macrometastases. Finally, we identified *Cd14*+ inflammatory neutrophils which were associated with TLR-NFκB inflammation signature gene expression and the CD14+ ‘activated’ MDSC module (**Fig. 4B**, bottom) and were proportionally enriched in the pre-metastatic niche (**Fig. 4B**, top right), as was observed in *Cd14*+ inflammatory AMs.

The co-detection of *Cd14*+ TLR-NFκB inflammatory neutrophils and AMs in the pre-metastatic niche led to the question of whether this transcriptional program reflects a broader mechanism of pre-metastatic niche development that can be detected in other immune cell types. To this end, we used NMF to identify components which were enriched in inflammatory neutrophils and AMs and assessed whether other myeloid subpopulations enacted these programs. This analysis identified a single component (NMF30) that was associated with *Cd14*+ inflammatory neutrophils and AMs (**Fig. S6A**) as well as subpopulations of monocytes and IMs (**Fig. 5A**). These IM and monocyte subpopulations reflect a coherent *Cd14* and TLR-NFκB-inducing phenomenon that is shared amongst lung myeloid cells, as inflammatory IMs and monocytes expressed high levels of TLR-NFκB inflammation signature genes (**Fig. 5B**) and were proportionally-enriched in the pre-metastatic niche (**Fig. 5C**). Moreover, NMF30 correlated with the CD14+ ‘activated’ MDSC signature (**Fig. 5A**, right; **Fig. 5D**) and was associated with inflammatory pathways such as TNF signaling via NFκB using GSEA (**Fig. 5E**), as was observed for the AM-specific NMF25. Notably, NMF30 did not exhibit enrichment amongst DCs nor did any DC subtypes co-express *Cd14*, *Cxcl2*, and *Nlrp3* in our data (**Fig. S6B**).

**Figure 5.**
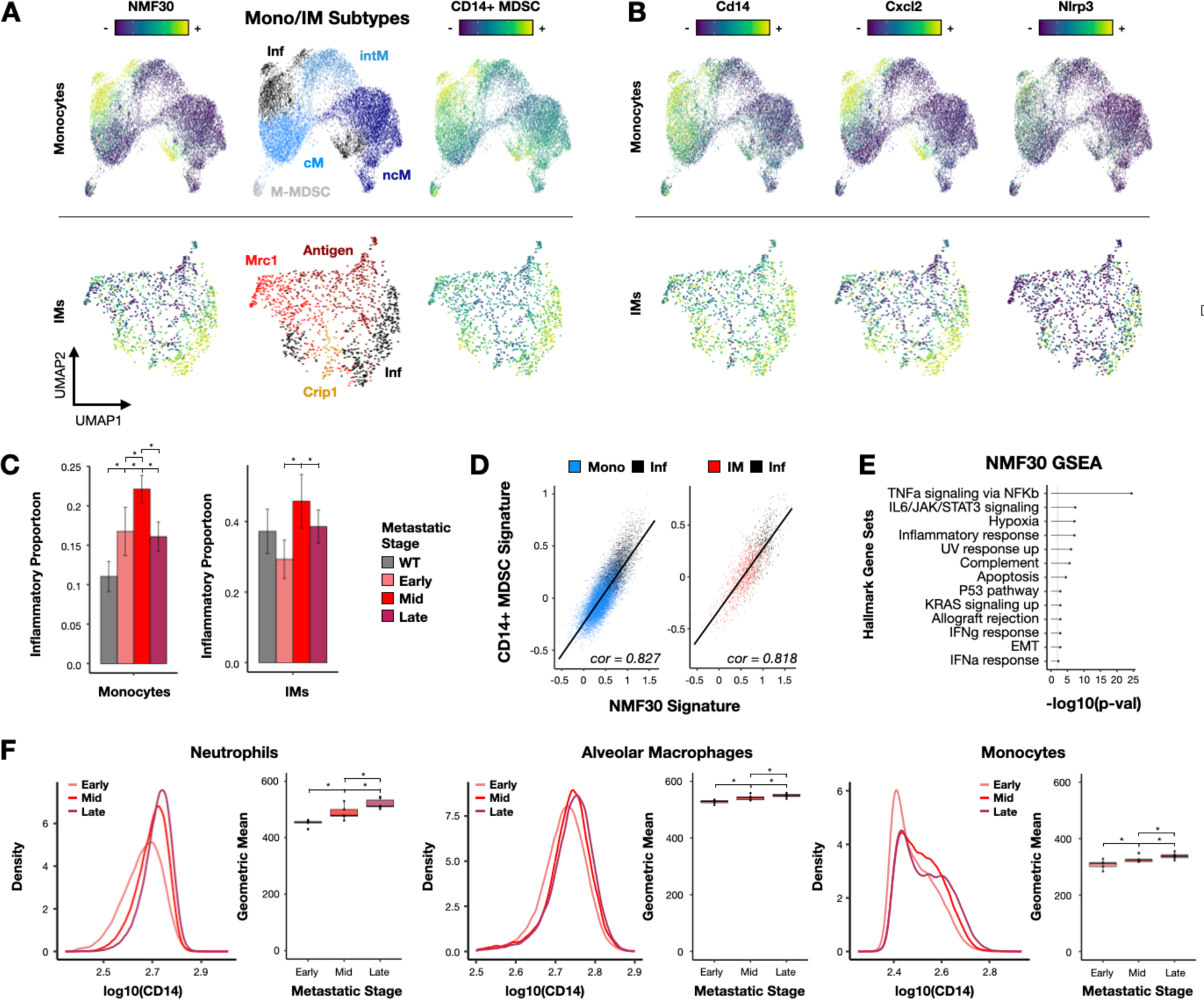
TLR-NFκB inflammation signature is present in BM-derived and tissue-resident myeloid cells and correlates with pre-metastatic niche formation. (A) Visualization of monocyte (top), and IM (bottom) gene expression space colored by NMF30 score (left), subtype (middle), or CD14+ ‘activated’ MDSC signature scores (right). (B) Visualization of monocyte (top) and IM (bottom) gene expression space colored by TLR-NFκB inflammation signature gene expression. (C) Bar charts describing shifts in inflammatory monocyte (left) and IM (right) subpopulations during metastatic progression. Data represented as mean ± s.e.m. * corresponds to p < 0.01 in two-proportion z-tests. (D) Scatter plot illustrating correlation between CD14+ ‘activated’ MDSC and NMF30 module scores in monocytes (left) and IMs (right). (E) Lollipop plot depicting Hallmark gene sets that are significantly enriched amongst genes comprising the NMF30 transcriptional signature. (F) Flow cytometry analysis of CD14 abundance on neutrophils (left), AMs (middle), and monocytes (right) binned by metastatic stage summarized using box plots. Dotted lines correspond to CD14 geometric mean. * corresponds to p < 0.01 in Wilcoxon rank-sum test.

Since TLR-NFκB inflammation in the pre-metastatic niche has not been described in these myeloid cell types, we used flow cytometry to measure changes in membrane CD14 protein abundance in these cells during metastatic progression. This analysis was aligned with our scRNA-seq data, as membrane CD14 abundance was up-regulated in neutrophils, AMs, and monocytes over time (**Fig. 5F; Fig. S7**). Notably, membrane CD14 protein abundance increased consistently between metastatic stages – differing from the transient increase observed in our scRNA-seq data – perhaps due to co-expression of the NMF30-associated gene Ccrl2, which increases CD14 membrane stabilization.^63^ Altogether, these analyses identify previously-unreported and metastasis-associated shifts in DC and neutrophil cell state and population structure, as well extend our understanding of established mechanisms of pro-metastatic immune remodeling^17–21^ by expanding the list of cell types known to engage in TLR-NFκB inflammation in the pre-metastatic niche. Moreover, similarities between inflammatory lung myeloid cells and the reported ‘activated’ MDSC signature suggest that local MDSC ‘activation’ in the primary tumor may in fact represent a coherent transcriptional program that can be elicited systemically in a wide range of myeloid populations to promote tumor progression and metastasis.

### Lymphocytes respond and contribute to an inflammatory and immunosuppressive PyMT lung metastatic microenvironment

Lymphocytes have diverse, context-dependent influences on metastatic progression. For instance, inflammation-induced NK and CD8 T cell cytotoxic responses are generally anti-metastatic, while immunosuppressive Tregs, regulatory B cells (Bregs), and ‘tumor-exposed’ NK cells can play pro-metastatic roles.^28, 31, 64–66^ Moreover, prior work has demonstrated that inflammation and immunosuppression are critical features of the metastatic microenvironment, as tumors hijack immune mobilization mechanisms while circumventing the cytotoxic responses normally associated with inflammation.^19, 20, 67^ Analysis of lung lymphocytes in our longitudinal scRNA-seq atlas revealed signatures that provide new molecular details into this phenomenon.

In NK cells, we identified immunomodulatory and cytotoxic subtypes that were proportionally biased towards cytotoxic NK cells over time (**Fig. 6A; Fig. S8A; Table S5**). Although cytotoxic NK cells are known to accumulate at inflammatory sites,^68^ this result was surprising because cytotoxic NK cells are thought to be anti-metastatic^64^ and NK cells in the PyMT primary tumor microenvironment are reprogrammed into an immunomodulatory and pro-metastatic state.^65^ We validated this result using flow cytometry (**Fig. S9**) which demonstrated that CD11b-high cytotoxic NK cell proportions indeed increased over time relative to CD27-high immunomodulatory NK cells (**Fig. 6B**). In T cells, we also observed that CD8 T cell differentiation increased over time (**Fig. 6C; Fig. S8B; Table S5**), which further demonstrates that lymphocytes respond to inflammatory cues in the metastatic niche.

**Figure 6.**
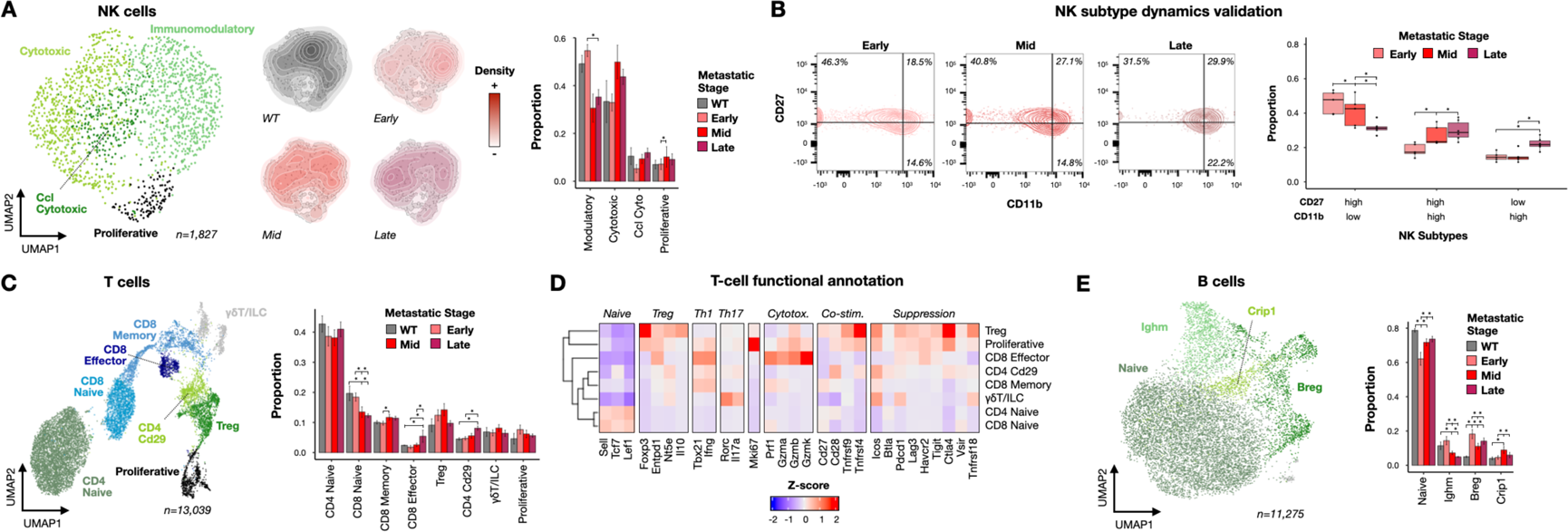
Lymphocyte subtype characterization reveals details of the inflammatory and immunosuppressive metastatic microenvironment. (A) Visualization of NK cell gene expression space colored by subtype (left) or metastatic stage densities (middle) with bar charts describing shifts in NK subtype proportions during metastatic progression (right). Proportional data represented as mean ± s.e.m. * corresponds to p < 0.01 in two-proportion z-tests. (B) Flow cytometry analysis of lung NK cells during metastatic progression. Data presented as CD11b x CD27 contour plots binned by metastatic stage and summarized using box plots. * corresponds to p < 0.01 in two-proportion z-tests. (C) Visualization of T cell gene expression space colored by subtype (left) with bar charts describing shifts in T cell subtype proportions during metastatic progression (right). Proportional data represented as mean ± s.e.m. * corresponds to p < 0.01 in two-proportion z-tests. (D) Z-score heatmap of T cell functional markers in each T cell subtype. (E) B cell analyses as presented in Fig. 6C.

Alongside these inflammatory responses, we also observed metastasis-associated signatures of lymphocyte immunosuppression. For instance, functional annotation of T cell subtypes (as performed previously^69^) revealed that the expression of cytotoxicity genes in *Cd29*+ cytotoxic CD4 T cells^70^ was suppressed (**Fig. 6D**), and that the proportion of immunosuppressive Tregs were enriched in the proliferative T cell compartment (**Fig. S8C**) and marginally increased in mid-stage lungs (**Fig. 6C**). Moreover, in B cells (**Fig. 6E; Fig. S8D; Table S5**) we observed that Breg frequencies were increased in all metastatic stages relative to WT mice, matching prior reports^31, 66^ and further demonstrating how T and B lymphocytes are biased towards immunosuppressive phenotypes. Considered collectively, these results demonstrate how distinct lymphocyte lineages participate in the inflammatory and immunosuppressive metastatic niche and suggest that lung- and primary tumor-associated NK cells are uniquely regulated during metastatic progression in PyMT mice.

### Intercellular communication network modeling predicts metastasis-associated immune signaling patterns

Our in-depth longitudinal characterization of lung immune cells during metastasis documented two key features of tumor-mediated immune remodeling: changes in immune cell state and population structure. However, metastatic progression is also associated with disease-specific patterns of intercellular communication.^31^ To assess how immune cell-cell signaling networks change during metastatic progression in PyMT mice, we used CellChat^71^ to predict active intercellular communication based on the expression of cognate receptor-ligand pairs at each metastatic stage.

To benchmark these predictions, we first assessed whether the predicted signaling networks for cytokines associated with *Cd14*+ TLR-NFκB inflammation – e.g., *Cxcl2*, *Tnf*, and *Il1b* (**Table S4**) – were distinct in pre-metastatic niche lungs, as would be expected for a pre-metastatic niche-linked transcriptional program. This analysis supported the accuracy of CellChat predictions, as *Cxcl2*-*Cxcr2* signaling primarily between AMs, IMs, and neutrophils (**Fig. 7A**) and *Tnf*-*Tnfrsf1a*/*Tnfrsf1b* signaling primarily between monocytes and AMs were predicted to be elevated in mid-stage lungs (**Fig. S10A**). Notably, *Il1b* signaling between immune cells was not predicted to be active as expression of the cognate receptor, *Il1r1*, was not detected (**Fig. S10B**, left). Instead, *Cd14*+ inflammatory neutrophils and AMs expressed the *Il1b* decoy receptor, *Il1rn* (**Fig. S10B**, right), suggesting that these cells cannot respond to *Il1b*.^72^

**Figure 7:**
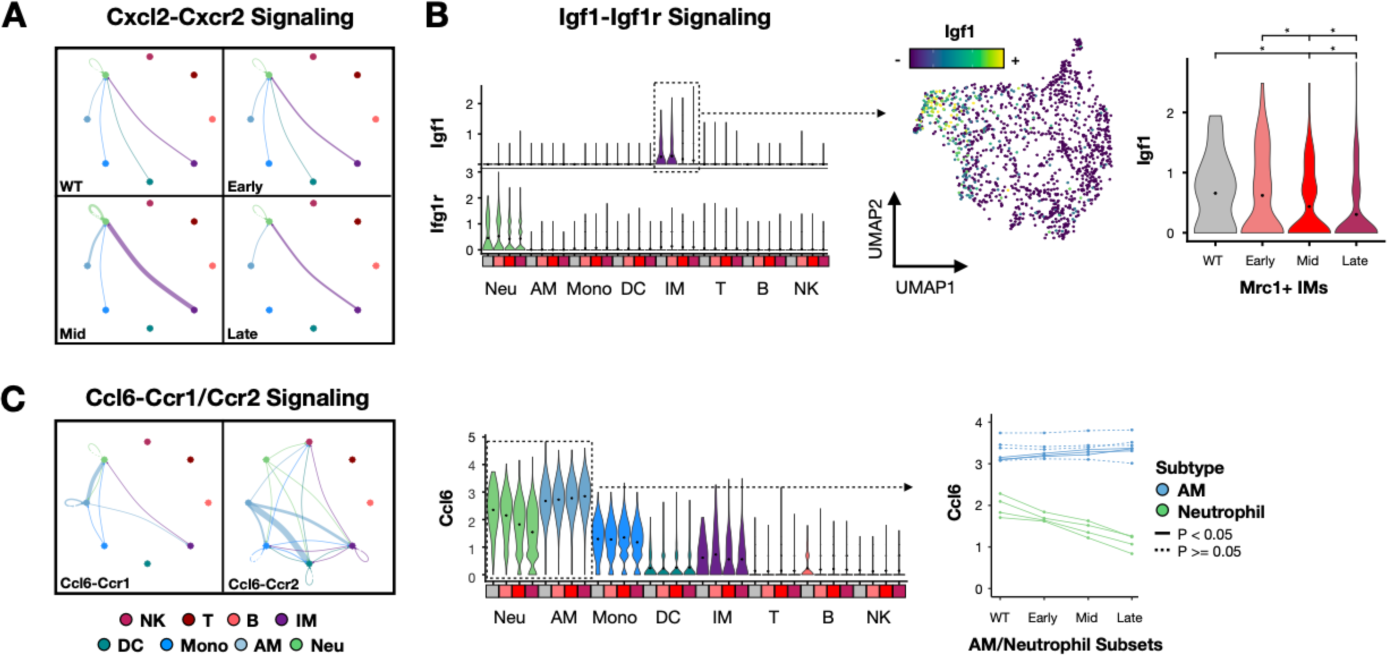
Intercellular communication modeling reveals metastasis-associated changes in immune cell signaling network. (A) Weighted network graph of *Cxcl2*-*Cxcr2* signaling interactions binned by metastatic stage. Nodes are colored by cell type, edges are weighted by signaling probability and colored by sender cell type. (B) Violin plots showing *Igf1* and *Igf1r* expression in cell types binned by metastatic stage (left) with IM gene expression space colored by *Igf1* expression (middle) and violin plot showing *Igf1* expression in *Mrc1*+ IMs binned by metastatic stage (right). Black dots on violin plots denote mean expression. * corresponds to p < 0.01 in Wilcoxon rank-sum test. (C) Weighted network graph of *Ccl6*-*Ccr1* and *Ccl6*-*Ccr2* signaling interactions (left) with violin plots showing *Ccl6* expression in cell types binned by metastatic stage (middle) and a scatter plot summarizing changes in *Ccl6* expression amongst AM (blue) and neutrophil (green) subtypes binned by metastatic stage. Black dots on violin plots denote mean expression. Solid lines in the scatter plot denote p < 0.05 in Wilcoxon rank-sum test. (C) Violin plots showing *Il1b* and *Il1r1* expression in cell types binned by metastatic stage (left) with neutrophil and AM gene expression spaces colored by Il1rn expression (middle) and violin plots showing Il1rn expression in inflammatory and non-inflammatory neutrophil and AM subsets. Black dots on violin plots denote mean expression. * corresponds to p < 0.01 in Wilcoxon rank-sum test.

After successfully benchmarking our signaling predictions, we used CellChat to identify signaling pathways that were associated with particular metastatic stages (**Fig. S10C**). Among these predicted interactions, we identified an *Igf1*-*Igf1r* signaling circuit between IMs and neutrophils which was specifically present in WT and early-stage lungs (**Fig. 7B**, left). Among IMs, *Igf1* was expressed only by *Mrc1*+ IMs (**Fig. 7B**, middle) and decreased over time (**Fig. 7B**, right). Since the proportion of *Mrc1*+ IMs did not shift over time (**Fig. 2A**, right), this observation suggests that changes in *Mrc1*+ IM phenotype underlie the observed absence of *Igf1*-*Igf1r* signaling at later metastatic stages. Notably, lung metastases are reduced in *Igf1r* knock-out mice after tumor tail-vail injection^73^. Thus, IM-neutrophil *Igf1*-*Igf1r* signaling may play a previously-undescribed role in early pre-metastatic niche formation

Finally, we observed a predicted signaling network consisting of *Ccl6*-expressing AMs, IMs, neutrophils, and monocytes and either *Ccr1*-expressing neutrophils or *Ccr2*-expressing monocytes, DCs, IMs, and NK cells (**Fig. 7C**, left). Interestingly, *Ccl6* expression decreased in neutrophils and increased in AMs over time (**Fig. 7C**, middle). This observation was not explained by shifts in neutrophil and AM subtype proportions, as *Ccl6* expression was reduced in all neutrophil subtypes and elevated in most AM subtypes during metastatic progression (**Fig. 7C**, right). While a recent study has demonstrated how *Ccl6* production by eosinophils promotes metastasis,^74^ differential cell-type-specific regulation of *Ccl6* signaling in AMs and neutrophils during metastatic progression has not been described. Further studies aiming to uncover the mechanisms underlying the observed cell-type-specific regulation of *Ccl6* expression in neutrophils and AMs may yield further insights tumor-mediated immune remodeling during metastasis.

## DISCUSSION

Improving our understanding how immune cells in distant organ sites are remodeled by primary tumor-derived cues during metastatic progression has the potential to reveal novel mechanisms of pro-metastatic immune dysregulation and inspire the first generation of anti-metastatic immunotherapies. However, most prior preclinical studies of tumor-immune interactions during metastasis either focused on the primary tumor microenvironment, lacked sufficient assay sensitivity or temporal sampling resolution, or used experimental systems which do not model the fully intact immune system or metastatic cascade. To address this knowledge gap, we used MULTI-seq to construct a longitudinal scRNA-seq atlas of lung immune cells isolated from PyMT tumor-bearing mice spanning the full trajectory of metastatic progression. In-depth computational analyses of these data revealed many metastasis-associated immune gene expression signatures which were only possible to detect because of our fine-grained temporal sampling strategy.

For example, we detected a TLR-NFκB inflammatory signature in neutrophils, monocytes, IMs and AMs which specifically correlated with pre-metastatic niche formation. Although TLR-NFκB inflammation is known to organize the metastatic microenvironment, this phenomenon has been primarily understood to involve tumor-induced expression of endogenous TLR ligands which drives NFκB signaling and recruitment of immunosuppressive myeloid cells from the periphery.^17–21^ Detecting neutrophil, monocyte, AM, and IM engagement of TLR-NFκB inflammation thus expands our knowledge of which cell types participate in the phenomenon of pro-metastatic TLR-NFκB inflammation. Moreover, we observed that inflammatory lung myeloid cells in the pre-metastatic niche exhibited elevated expression of the TLR co-receptor, *Cd14*, which may facilitate their ability their activate TLR-NFκB signaling. The induction of *Cd14* expression and TLR-NFκB signaling mirrored reports of CD14+ ‘activated’ primary tumor MDSCs^50^ despite differences in cellular identity, ontology, and location, which suggests that primary tumor-derived cues induce a conserved myeloid gene expression program operates systemically and promotes both tumor progression and metastasis. Future efforts to pinpoint which tumor-immune signaling interactions drive *Cd14*+ TLR-NFκB inflammation will continue to improve our understanding of this mechanism of pro-metastatic immune remodeling and may facilitate anti-metastatic immunotherapy development.

Beyond myeloid TLR-NFκB inflammation, we also observed lymphocyte contributions to the inflammatory and immunosuppressive metastatic microenvironment which were highlighted by our observation that cytotoxic NK cell proportions were increased during metastatic progression. This result was surprising because cytotoxic NK cells are anti-metastatic^64^ and NK cells in the PyMT primary tumor microenvironment are reprogrammed into an immunomodulatory and pro-metastatic state,^65^ which suggests that NK cells are differentially regulated in PyMT primary tumors and lungs. Notably, cytotoxic NK cells were anecdotally shown to accumulate in metastasized lymph nodes isolated from melanoma patients,^75^ and NK cell activation is consistent with active inflammation^68^ and tumor cell downregulation of NK-inhibitory MHC-I^76^ in the metastatic niche. However, how metastatic tumors circumvent NK cell cytotoxicity in this context remains unclear. One possibility is that NK cell cytotoxicity is suppressed in an orthogonal fashion to NK cell activation. Indeed, while analysis of cytotoxic NK cell transcriptional profiles did not reveal any stage-specific differences, our flow cytometry data showed that CD11b+ mid-stage NK cells expressed elevated levels of the immunomodulation marker CD27 relative to late-stage NK cells. Thus, deeper scRNA-seq profiling of NK cells during metastatic progression with paired functional, phenotypic, and spatial transcriptomic assays may provide new details on how metastatic tumor cells circumvent NK cell cytotoxicity. Perturbing these NK cell-suppressive pathways may also prove fruitful for anti-metastatic immunotherapy development.

Finally, we described a variety of metastasis-associated immune gene expression signatures in this study that are aligned with known biological principles but must be experimentally validated. For example, detection of wound healing-associated *Crip1*+ IMs and non-cytotoxic *Cd29*+ CD4 T cells are consistent with dynamic reorganization of an immunosuppressive lung microenvironment but have not been reported in previous studies. Similarly, decreases in cDC1 and B cell proportions over time are in line with systemic suppression of adaptive immune responses during metastasis, but causal connections between these observations and metastatic progression must be empirically tested. Finally, while IGF1 and CCL6 signaling have both been associated with lung metastatic progression,^73, 74^ whether the predicted suppression of IM-neutrophil *Igf1*-*Igf1r* signaling and cell-type-specific regulation of *Ccl6* expression that we observed during metastatic progression contribute to disease progression must be explicitly assessed. Causally linking IGF1 and CCL6 signaling to lung metastatic niche development would position these two pathways as prime candidates for anti-metastatic immunotherapy development.

Considered collectively, our study identifies literature-supported and novel changes in immune cell gene expression profiles, population structure, and cell-cell signaling interactions which correlate with metastatic progression. These observed changes provide new insights into established organizing mechanisms in the metastatic microenvironment, identify candidate pathways for anti-metastatic immunotherapy development, and put forth testable hypotheses to improve our understanding of tumor-mediated immune remodeling during metastasis. We anticipate that the longitudinal scRNA-seq atlas presented here will serve as a foundational resource for the broader community as PyMT mice are widely used in cancer immunology research.

### Limitations of the study

There are three key limitations of this study. First, we cannot comment on how non-immune stromal cells such as fibroblasts, epithelial cells, and vascular endothelial cells are remodeled during metastatic progression. Since these cell populations are also known to promote metastasis in certain contexts,^21, 77–79^ future longitudinal scRNA-seq analyses of non-immune stromal cells may provide more insight into how the metastatic niche is organized. Second, while we identify a variety of metastasis-associated immunological signatures, we do not establish causal links to metastatic progression for any of the signatures described in this study. Follow-up studies which specifically interrogate whether blocking *Igf1*-*Igf1r* IM-neutrophil signaling, myeloid TLR-NFκB inflammation, and/or *Cd14*-inducing primary tumor-derived cues using traditional molecular biology and immunology approaches are needed to assess precisely how these pathways regulate metastatic progression. Finally, whether the immunological signatures we observe in the PyMT lung metastatic nice are generalizable across different primary tumor types and metastasized organs remains unclear. Expanding our longitudinal scRNA-seq experimental design to different metastatic cancer mouse models and primary patient samples may yield further insights into the organizing principles of tumor-mediated immune remodeling and bring anti-metastatic immunotherapies one step closer to reality.

## ACKNOWLEDGEMENTS

This research was funded in part through NCI U01 CA260852 and the Parker Institute for Cancer Immunotherapy. C.S.M. is a Cancer Research Institute Irvington Fellow supported by the Cancer Research Institute (CRI Award #4134) and a Parker Institute for Cancer Immunotherapy Scholar. A.T.S. was supported by a Career Award for Medical Scientists from the Burroughs Wellcome Fund, a Lloyd J. Old STAR Award from the Cancer Research Institute, a Pew-Stewart Scholars for Cancer Research Award, and the Donald and Delia Baxter Foundation. We thank Zev Gartner and Danny Conrad for providing MULTI-seq reagents, Katalin Sandor for helping with optimizing tissue dissociation protocols, and Kamir Hiam-Galvez for intellectual discussions.

## AUTHOR CONTRIBUTIONS

C.S.M. conceived the project, performed experiments, analyzed the data, and wrote the manuscript. Z.M. performed experiments. N.E.R. and J.W. wrote the manuscript. A.T.S. supervised the study and wrote the manuscript.

## RESOURCE AVAILABILITY

### Lead contact

Further information and requests for resources should be directed to and will be fulfilled Chris McGinnis and Ansuman Satpathy.

### Materials availability

This study did not generate new reagents.

### Data and code availability

Raw mapped count matrices, pertinent metadata, and FASTQs can be found in the National Center for Biotechnology information Gene Expression Omibus with accession number GSEXXXXXX. R scripts used for analyzing data and generating all manuscript figures are deposited on GitHub: github.com/chris-mcginnis-ucsf/pymt_atlas

## EXPERIMENTAL METHODS

### Animals

FVB/N-Tg(MMTV-PyVT)634Mul/J mice were purchased from Jackson laboratories (strain #002374) and bred to produce hemizygous MMTV-PyMT females which were used for all experiments. Mice were housed in a pathogen-free facility. All experiments were performed according to protocols approved by the Institutional Animal Care and Use Committees of Stanford University (protocol number: 33814).

### Lung dissociation and cryopreservation

Lung single-cell suspensions were prepared for cryopreservation using the following workflow. Mice were first euthanized and lungs were surgically dissected and placed in ice-cold PBS. Each lung was then fragmented using surgical scissors and transferred to 5 mL of dissociation media (1 lung per dissociation reaction) in gentleMACS™ C Tubes (Miltenyi Biotec, 130-093-237). Dissociation media contained 0.5 U/mL dispase (Stemcell Technologies, 07923), 0.1 mg/mL DNase I (Sigma-Aldrich, 11284932001), 490 µL/mL Advanced DMEM/F-12 (Fisher Scientific, 12-634-028), and 2 mg/mL collagenase P (Sigma-Aldrich, 11249002001). Lungs were then dissociated for 20 minutes using a custom GentleMACS protocols before filtering through a 70 µm Macs SmartStrainer (Miltenyi Biotec, 130-110-916) into Advanced DMEM/F-12 supplemented with 2% FBS (DMEM-FBS). Tissue fragments remaining on the filter were then transferred with forceps to 5 mL of fresh dissociation media and dissociated again for 20 minutes before filtering to combine with the previous filtrate. Cells were then pelleted (400xg, 4°C, 4 minutes), washed once with 1 mL of DMEM-FBS, and resuspended in 1 mL of Recovery™ Cell Culture Freezing Medium (Fisher Scientific, 12-648-010). Cells in freezing medium were kept at −80°C for 24-48 hours before transferring to LN2 for long-term storage. Left lungs were designated for scRNA-seq analysis while right lungs were used for matched flow cytometry validation experiments.

### scRNA-seq sample preparation

Cryopreserved left lung single-cell suspensions were prepared for scRNA-seq using the following workflow. First, samples were thawed in a 37°C water-bath and transferred drop-wise to 12 mL of DMEM-FBS. Cells were then pelleted (400xg, 4°C, 4 minutes) and resuspended in 500 µL of DMEM-FBS and counted. 5×10^5^ total cells from each sample were then aliquoted into distinct wells of a round-bottom 96-well plate and washed once with 250 µL of PBS. Cells were then resuspended in 80 µL of PBS and stained for 5 minutes on ice with 10 µL of MULTI-seq anchor lipid-modified oligonucleotide (LMO) pre-hybridized to a unique MULTI-seq barcode (2 µM stock, 200 nM labeling concentration).^16^ After staining for another 5 minutes on ice with 10 µL of MULTI-seq co-anchor LMO (2 µM stock), the cells were diluted with 150 µL of 1% BSA in PBS (PBS-BSA) and pooled into a 15 mL conical tube. The sample pool was washed once with 10 mL of PBS-BSA and resuspended in 150 µL of Zombie NIR viability dye (1:500 in PBS; BioLegend 423105). After 15 minutes on ice, the cells were diluted with 5 mL of 2% FBS in PBS (FACS buffer), pelleted, and resuspended in 150 µL of Fc-block (1:200 in FACS buffer; Tonbo, 700161U500). After 5 minutes on ice, the cells were diluted with 5 mL of FACS buffer, pelleted, and resuspended in 150 µL of an antibody cocktail containing 1:200 anti-Ter119 (FITC; Thermo Scientific, 11-5921-82), 1:50 anti-CD31 (FITC; Thermo Scientific, 11-0311-82), and 1:40 anti-CD45 (violetFluor™ 450; Tonbo, 75-0451-U100) mouse monoclonal antibodies in FACS buffer. After 45 minutes on ice, the cells were diluted with 5 mL of FACS buffer, washed once with 5 mL of FACS buffer, and filtered through a 70 µm Macs SmartStrainer prior to FACS enrichment for CD45+ Ter119-CD31-live immune cells (excluding red blood cells and endothelial cells) using a BD FACSAria II instrument. After FACS, cells were counted, the concentration was adjusted to 1×10^6^ cells/mL, and 43.2 µL of the cell suspension was ‘super-loaded’ into seven microfluidic lanes of a 10x Genomics 3’ scRNA-seq V3.1 chip. Each metastatic timepoint (weeks 6-14) and 9-week old WT controls were represented with three biological replicates along with one 15-week old PyMT mouse.

### scRNA-seq library preparation and next generation sequencing

scRNA-seq libraries were prepared according to manufacturer’s recommendations (10x Genomics). MULTI-seq libraries were prepared as described previously.^16^ scRNA-seq and MULTI-seq libraries were pooled and sequenced using NovaSeq 6000 S4 flow cells to an average sequencing depth of 35,000 reads/cell and 3,000 reads/cell, respectively.

### Flow cytometry

Cryopreserved right lung single-cell suspensions were prepared for flow cytometry analysis using the following workflow. First, samples were thawed in a 37°C water-bath and transferred drop-wise to 9 mL of 10% FBS in PBS. Cells were then pelleted (400xg, 4°C, 4 minutes) and resuspended in 1 mL of 2% FBS in PBS (FACS buffer). Cells were then pelleted and resuspended in 100 µL of Zombie Green viability dye (1:500 in PBS; BioLegend, 423111). After 15 minutes on ice, the cells were diluted with 1 mL of FACS buffer, pelleted, and resuspended in 100 µL of Fc-block (1:200 in FACS buffer). After 5 minutes on ice, the cells were diluted with 1 mL of FACS buffer, pelleted, and resuspended in 100 µL of antibody cocktail in FACS buffer. The antibody panels for NK cell and myeloid analyses were designed as described previously.^80–82^ The compositions of the antibody cocktails were as follows:

| NK cocktail | Fluorophore | Dilution | Vendor | Catalogue # |
| --- | --- | --- | --- | --- |
| Ter119 | FITC | 1:200 | Thermo Scientific | 11-5921-82 |
| CD31 | FITC | 1:200 | Thermo Scientific | 11-0311-82 |
| CD3 | FITC | 1:200 | Thermo Scientific | 11-0032-82 |
| CD19 | FITC | 1:200 | Thermo Scientific | 11-0193-81 |
| THY1.1 | PE | 1:333 | Thermo Scientific | 12-0900-81 |
| CD27 | APC | 1:80 | BioLegend | 124211 |
| CD11b | PerCP-Cy5.5 | 1:80 | Thermo Scientific | 45-0112-80 |
| KLRG1 | eFluor 506 | 1:40 | Thermo Scientific | 69-5893-82 |
| NKp46 | PE-Cy7 | 1:20 | Thermo Scientific | 25-3351-80 |
| CD45 | APC-Cy7 | 1:20 | Thermo Scientific | A15395 |
| CD49b | eFluor 450 | 1:10 | Thermo Scientific | 48-5971-80 |

| Myeloid cocktail | Fluorophore | Dilution | Vendor | Catalogue # |
| --- | --- | --- | --- | --- |
| Ter119 | FITC | 1:200 | Thermo Scientific | 11-5921-82 |
| CD31 | FITC | 1:200 | Thermo Scientific | 11-0311-82 |
| SIGLEC-F | PE-Cy7 | 1:160 | Thermo Scientific | 25-1702-82 |
| CD11b | PerCP-Cy5.5 | 1:80 | Thermo Scientific | 45-0112-80 |
| MHC-II | BV421 | 1:80 | Thermo Scientific | 404-5321-80 |
| Ly6G | APC | 1:40 | Tonbo | 20-1276-U025 |
| CD64 | BV605 | 1:40 | BioLegend | 139323 |
| CD45 | APC-Cy7 | 1:20 | Thermo Scientific | A15395 |
| CD14 | PE | 1:20 | Thermo Scientific | 12-0141-81 |

After 30 minutes in antibody cocktails on ice, the cells were diluted with 1 mL of FACS buffer, pelleted, and washed twice with 1 mL of FACS buffer prior to filtering and analysis using a BD FACSAria II instrument.

## COMPUTATIONAL METHODS

### scRNA-seq library pre-processing, quality-control, and MULTI-seq sample classification

scRNA-seq library FASTQs were pre-processed using Cell Ranger version 6.0.0 (10x Genomics) and aligned to the mm-10-3.0.0 reference transcriptome. Cell Ranger aggregate was used to perform read-depth normalization. Raw read depth normalized scRNA-seq count matrices were then read into R and parsed to exclude cell barcodes with fewer than 10^2.5^ total UMIs and genes with fewer than 5 counts across all cell barcodes. Parsed scRNA-seq data was then pre-processed using the standard Seurat workflow^83^ and clusters with low total UMIs and/or high proportion of mitochondrial transcripts (pMito) were excluded.

Cell barcodes passing the first quality-control workflow were then used to pre-process MULTI-seq barcode FASTQs using the ‘deMULTIplex’ R package.^16^ MULTI-seq sample classification was then performed independently for each scRNA-seq library, as described previously.^16^ After incorporating MULTI-seq sample classifications as metadata, clusters enriched with MULTI-seq-defined doublets and unclassified cells were removed prior to re-processing using the SCTransform workflow.^84^ These data were used for manual cell type annotation before the data was subsetted by cell type and re-processed using the SCTransform workflow. Manual subtype annotation was then performed and low-quality/doublet cell clusters missed during the initial quality-control workflows were removed.

### Non-negative matrix factorization (NMF)

NMF was performed as implemented in the ‘nmf’ function of the ‘RcppML’ R package.^85^ Raw gene expression matrices subsetted to include the top 3,000 variably-expressed genes (defined using the ‘VariableFeatures’ Seurat function) were used for all NMF runs. Sample factor matrices were used to visualize NMF component scores in gene expression space. Feature factor matrices were visually thresholded to identify NMF component genes before submitting for GSEA using hypeR.^47^

### Computing gene expression signature scores

Per-cell gene expression signature scores for NMF components and the CD14+ ‘activated’ MDSC signature^50^ were computed using the ‘AddModuleScore’ Seurat function. CD14+ ‘activated’ MDSC signature genes were parsed to include only genes present in the 3,000 variably-expressed genes prior to calculation of signature scores.

### Intercellular signaling predictions

Intercellular signaling interactions were predicted using CellChat^71^ first by splitting the full scRNA-seq atlas data according to metastatic stage (e.g., WT, early, mid, and late) and processing individual CellChat objects using the demonstrated workflow. Processed CellChat objects were then merged using the ‘mergeCellChat’ function, enabling comparative visualizations of weighted cell-cell signaling networks.

### Statistical tests

Statistically-significant shifts in proportions in both scRNA-seq and flow cytometry data were assessed using the two-proportion z-test^86, 87^ implemented in the ‘prop.test’ R function (p = NULL, alternative = “two.sided”, correct = T). Differentially-expressed genes between clusters in all datasets were defined using the Wilcoxon rank-sum test as implemented in the ‘FindAllMarkers’ Seurat function. Specific differential expression testing was performed using the Wilcoxon rank-sum test as implemented in the ‘FindMarkers’ Seurat function. For flow cytometry analysis, statistically-significant differences in geometric mean CD14 abundance were assessed using Wilcoxon rank-sum test as implemented in the ‘wilcox.test’ R function (alternative = “two.sided”).

