## Supplemental Figures and Tables for "The temporal progression of immune remodeling during metastasis"

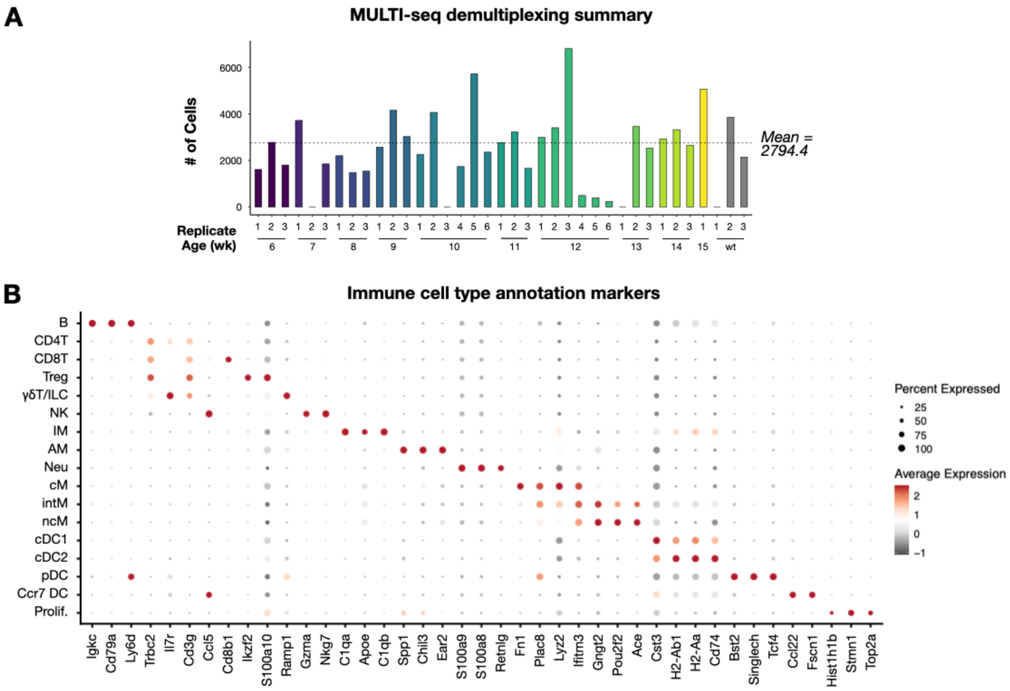


**Figure S1: Summaries of MULTI-seq classification and immune cell type annotation results.** Related to Figure 1.
(A) Bar charts describing the number of cells assigned to each sample following MULTI-seq demultiplexing.
(B) Dot plots for immune cell type annotation markers.

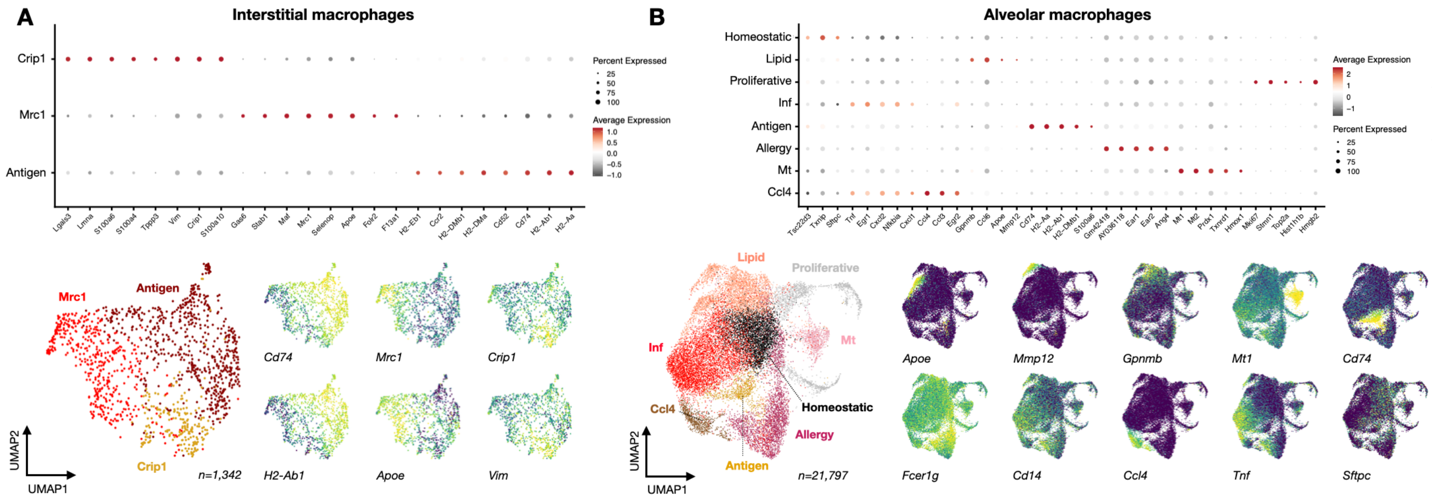


**Figure S2. Dot and feature plots for tissue-resident macrophage subtype annotation markers.** Related to Figure 2.
(A) Dot plots (top) and feature plots (bottom) for interstitial macrophage subtype annotation markers.
(B) Dot plots (top) and feature plots (bottom) for alveolar macrophage subtype annotation markers.

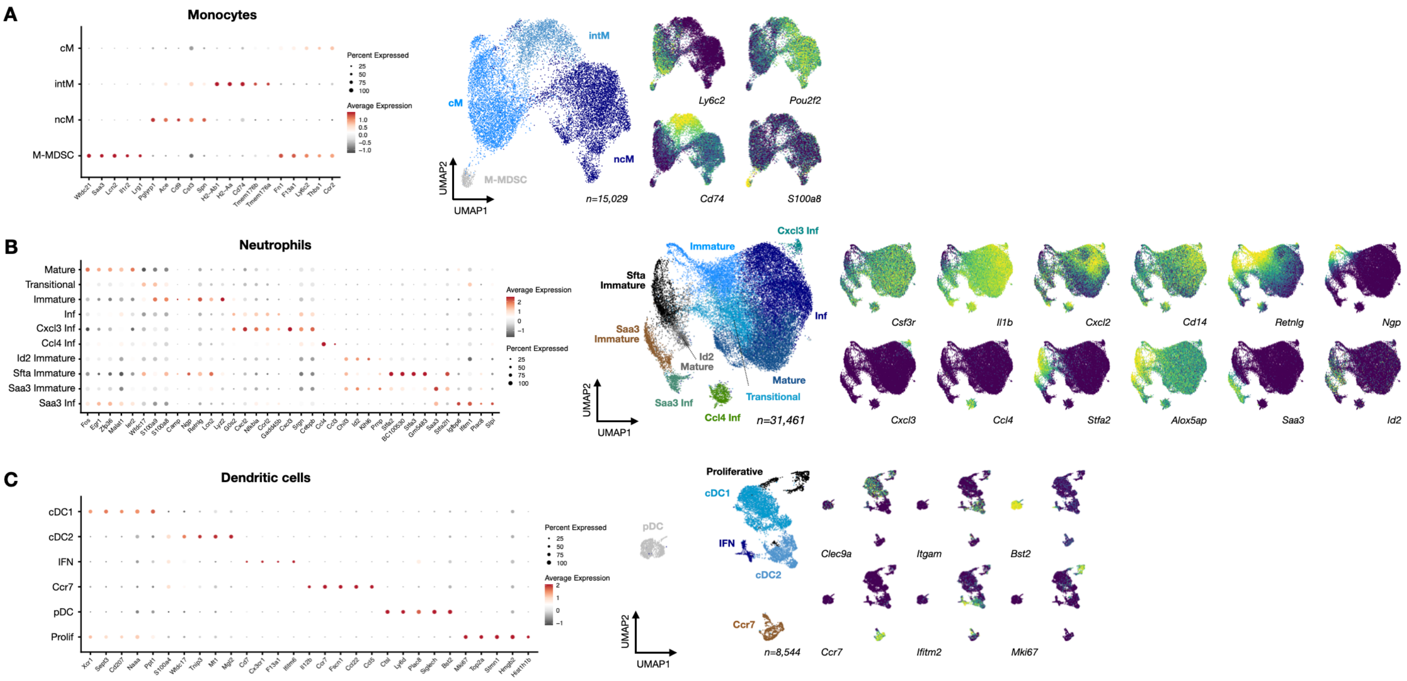


**Figure S3. Dot and feature plots for peripherally-derived myeloid subtype annotation markers.** Related to Figure 4.
(A) Dot plots (left) and feature plots (right) for monocyte subtype annotation markers.
(B) Dot plots (left) and feature plots (right) for neutrophil subtype annotation markers.
(C) Dot plots (left) and feature plots (right) for dendritic cell subtype annotation markers.

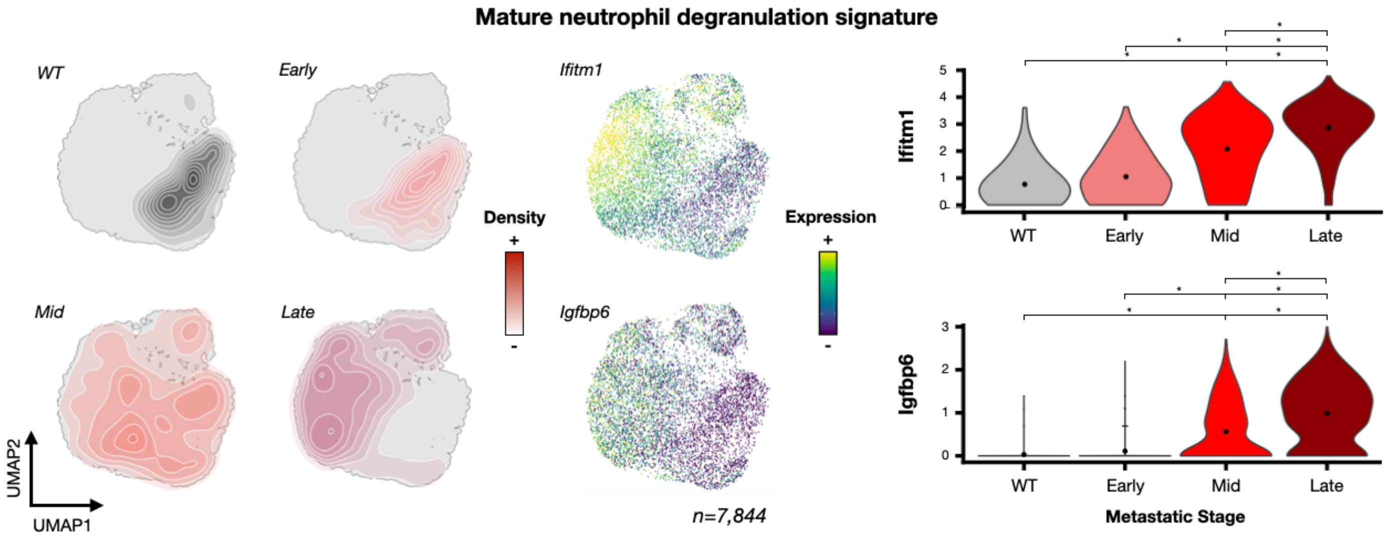


**Figure S4. Metastatic progression is associated with neutrophil degranulation signature in mature neutrophils**. Related to Figure 4. Visualization of metastatic stage densities in mature neutrophil gene expression space (left) with feature and violin plots (middle, right) of genes associated with neutrophil degranulation in vitro (e.g., *Igfbp6*) and in COVID-19 patients (e.g., *Ifitm1*). Black dot in violin plots denotes mean expression value. * corresponds to $p < 0.01$ in Wilcoxon rank-sum test.

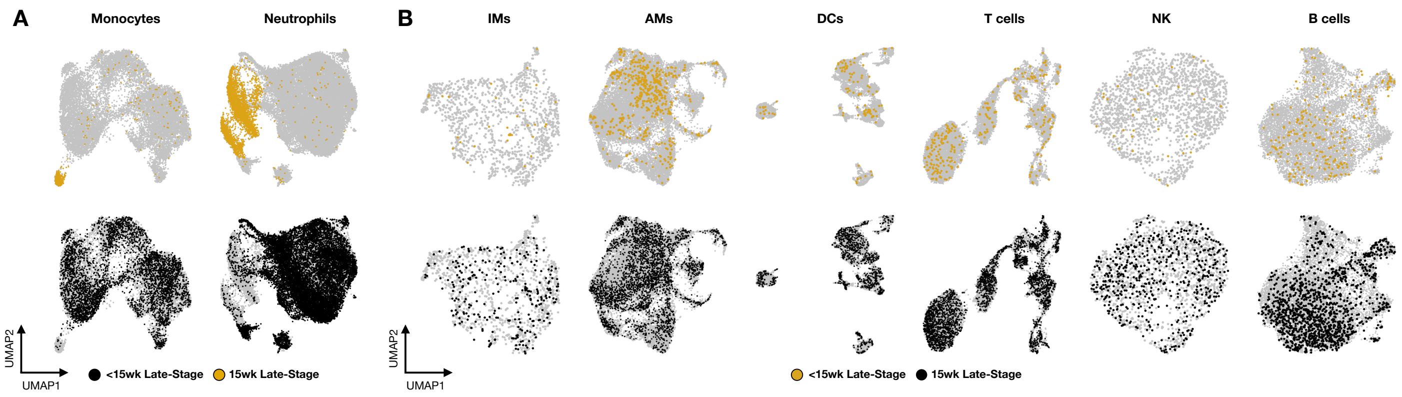


**Figure S5. Monocytes and neutrophils from 15-week PyMT mice are distinct from other late-stage metastasis lungs**. Related to Figure 4.
(A) Visualization of monocyte or neutrophil gene expression space highlighting cells from 15-week (top, gold) or <15-week late-stage metastasis lungs (bottom, maroon).
(B) Visualization of IM, AM, DC, T cell, NK, or B cell gene expression space highlighting cells from 15-week (top, gold) or <15-week late-stage metastasis lungs (bottom, black).

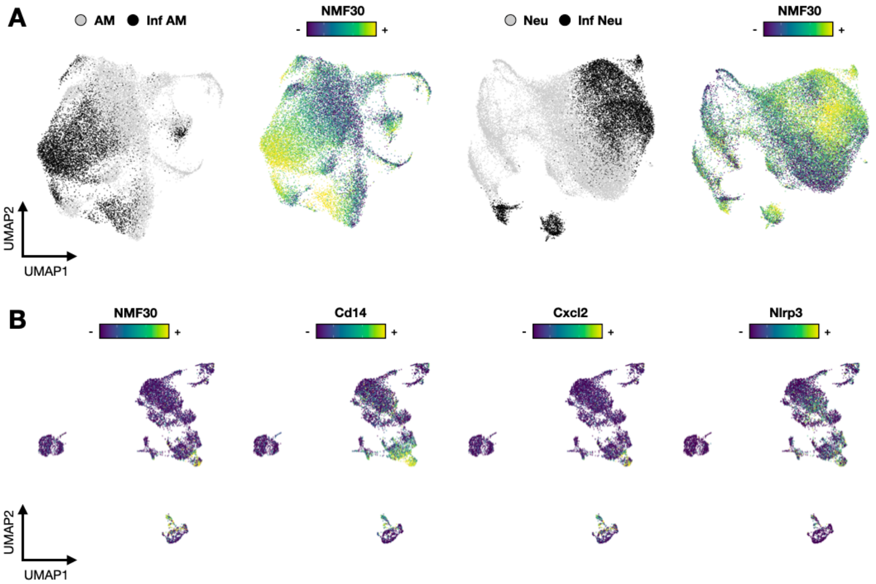


**Figure S6: NMF30 marks *Cd14*+ inflammatory neutrophils and AMs but not DCs.** Related to Figure 5.
(A) Visualization of AM (left) and neutrophil (right) gene expression space colored by NMF30 or inflammation status.
(B) Visualization of DC gene expression space colored by NMF30 or *Cd14*+ inflammation signature gene expression.

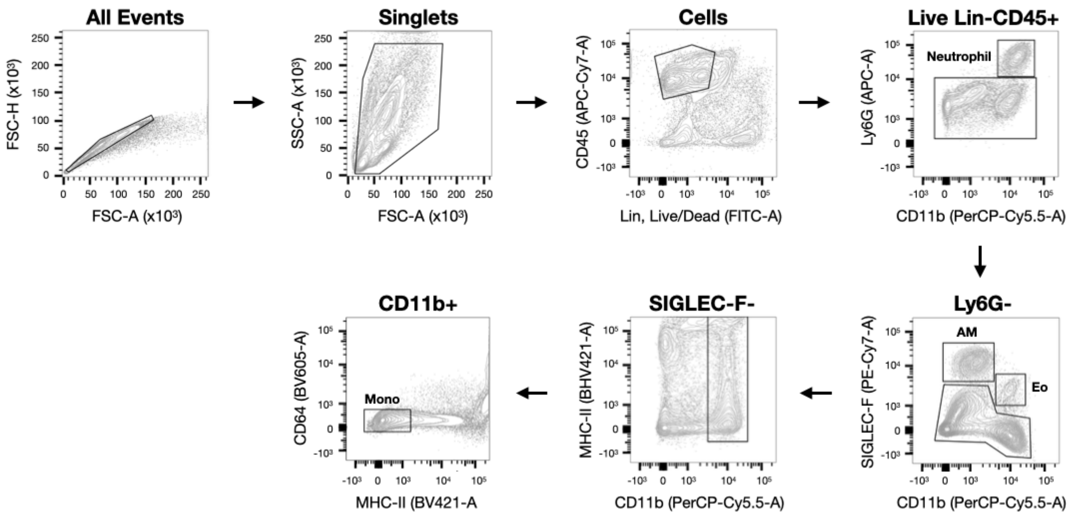


**Figure S7: Flow cytometry analysis of CD14 expression in neutrophils, AMs, and monocytes during metastatic progression.** Related to Figure 5.

Gating strategy to identify myeloid cells. Neutrophils were distinguished according to lack of CD31 and Ter119 expression (Lin gate) and expression of CD45, Ly6G, and CD11b. Amongst Ly6G- cells, AMs were distinguished according to low CD11b expression and high SIGLEC-F expression. Eosinophils (Eo) were also excluded at this step via high CD11b expression and intermediate SIGLEC-F expression. Amongst Ly6G-SIGLEC-F- myeloid cells, monocytes (mono) were then distinguished according to high CD11b expression and low MHC-II and CD64 expression.

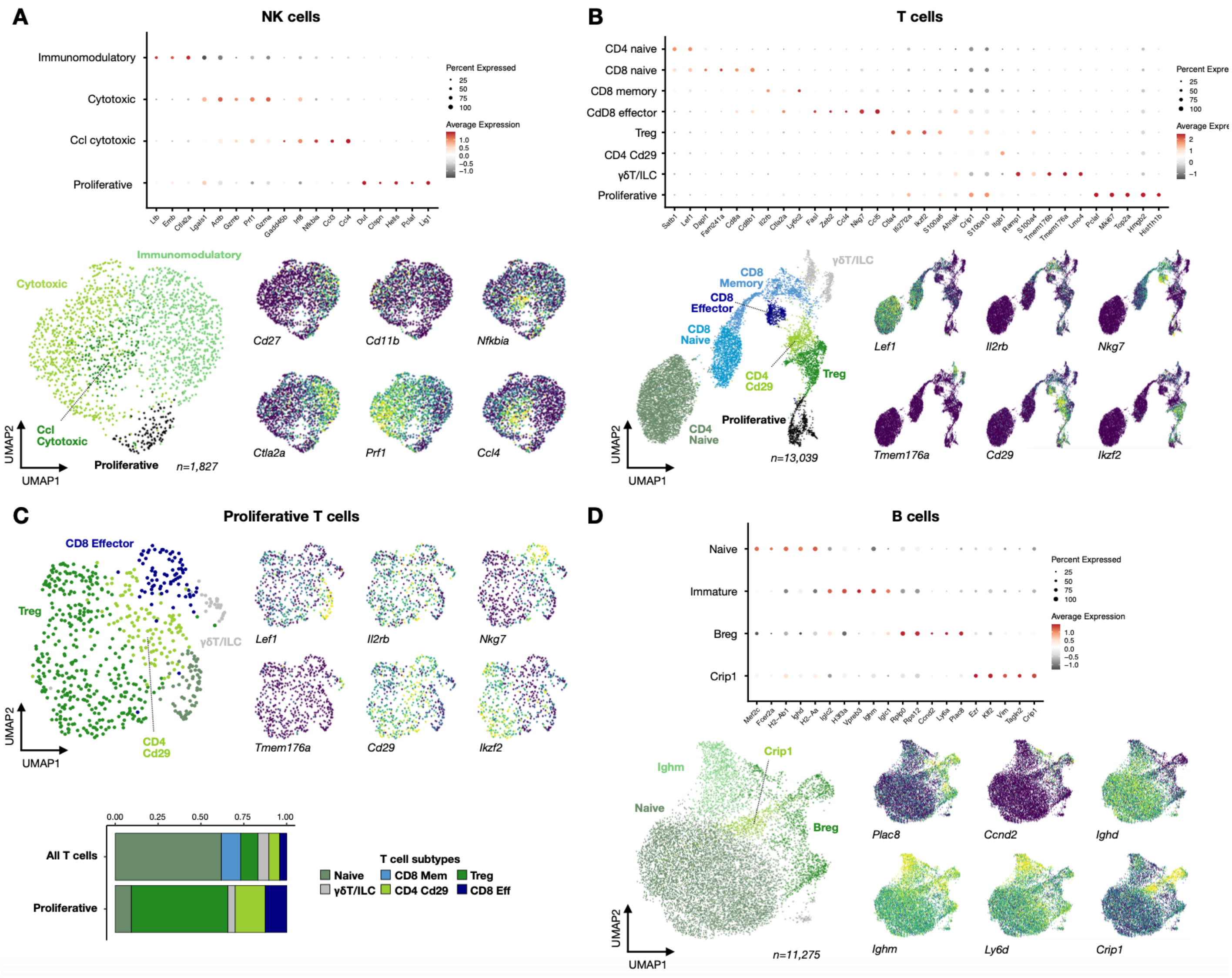


**Figure S8. Dot and feature plots for lymphoid subtype annotation markers.** Related to Figure 6.
(A) Dot plots (top) and feature plots (bottom) for NK cell subtype annotation markers.
(B) Dot plots (top) and feature plots (bottom) for T cell and ILC subtype annotation markers.
(C) Feature plots for proliferative T cell subtypes (top) with bar charts showing subtype proportions amongst proliferative and non-proliferative T cells.
(D) Dot plots (top) and feature plots (bottom) for B cell subtype annotation markers.

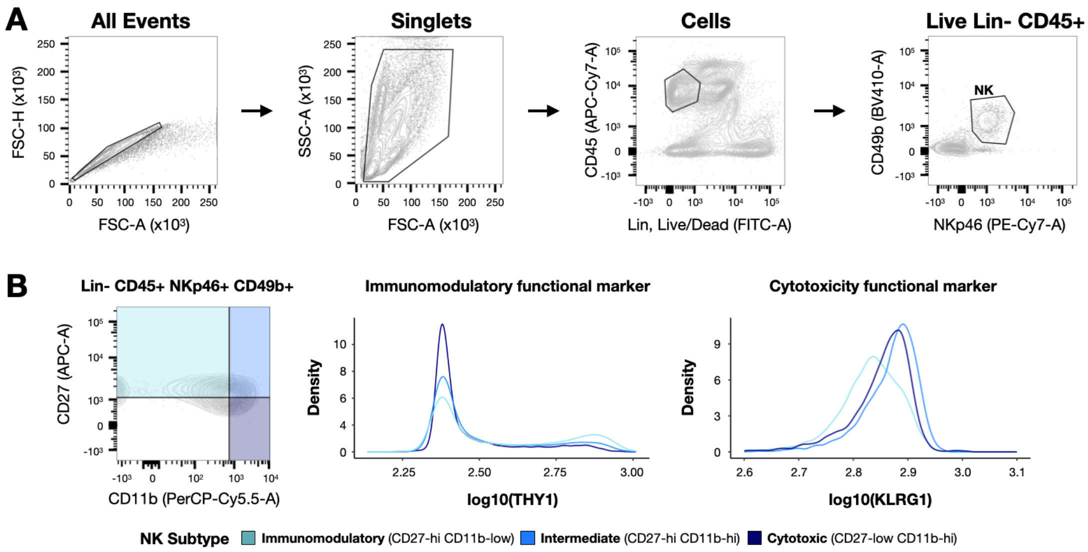


**Figure S9: Flow cytometry analysis of natural killer cell subtypes during metastatic progression.** Related to Figure 6.
(A) Gating strategy to identify NK cells. Live NK cells were distinguished were distinguished from other lung immune cells according to lack of CD3, CD19, Ter119, and CD31 expression (Lin gate) and expression of CD45, CD49b, and NKp46. Scatter plots three representative samples from early, mid, and late-stage lungs.
(B) Gating strategy to assess immunomodulatory and cytotoxic NK cell subtypes using quadrant gating of CD27 and CD11b abundance (left) with distributions of immunomodulatory (THY1) and cytotoxicity (KLRG1) functional markers binned by CD27/CD11b gate. THY1 abundance is highest in the immunomodulatory gate (light blue) while KLRG1 is highest in the intermediate (blue) and cytotoxic (dark blue) gates.

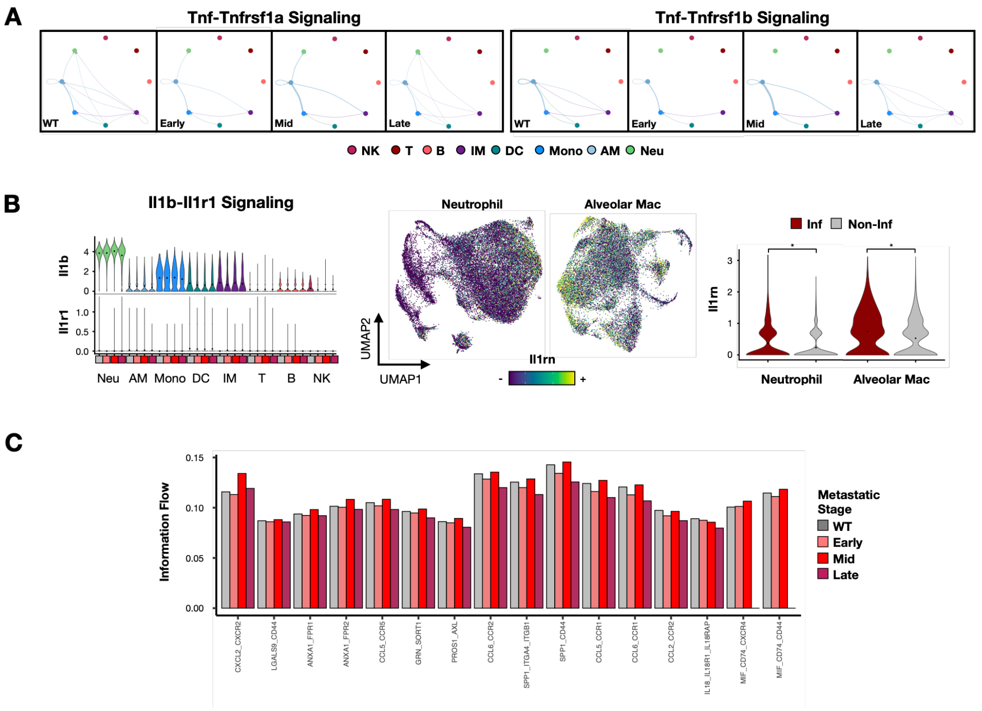


**Figure S10: Interrogating cell-cell communication predictions associated with the TLR-NFκB inflammatory signature.** Related to Figure 7.

(A) Weighted network graph of Tnf-Tnfrsf1a (left) and Tnf-Tnfrsf1b (right) signaling interactions split by metastatic stage. Nodes are colored by cell type, edges are weighted by signaling probability and colored by sender cell type

(B) Violin plots showing Il1b and Il1r1 expression in cell types binned by metastatic stage (left) with neutrophil and AM gene expression spaces colored by Il1rn expression (middle) and violin plots showing Il1rn expression in inflammatory and non-inflammatory neutrophil and AM subsets. Black dots on violin plots denote mean expression. * corresponds to $p < 0.01$ in Wilcoxon rank-sum test.

(C) Bar charts describing information flow during each metastatic stage for the top 16 predicted signaling interactions ranked by statistical significance.

| Table S1: Immune cell type markers and population structure dynamics (Related to Figure 1) | | |
|---|---|---|
| B | Igkc, Cd79a, Ly6d, Ebf1, Ighm, Cd79b, Iglc2, Ighd, Ms4a1, Iglc3 | Consistent decrease |
| CD4T | Trbc2, Il7r, Cd3g, Cd3d, Lef1, Trbc1, Trac, Cd3e, Ets1, Tmsb10 | Early decrease then stable |
| CD8T | Ccl5, Trbc2, Cd8b1, Cd3g, Cd3d, Nkg7, Cd3e, Trbc1, Tmsb10, Trac | Early decrease then stable |
| Treg | Ikzf2, Trbc2, S100a10, Ctla4, Ifi27l2a, Cd3g, Trbc1, Tnfrsf4, S100a4, Cd3e | Consistent decrease |
| γδT/ILC | Il7r, Ramp1, Cd3g, Cxcr6, Trbc1, S100a4, Rora, Trbc2, Tmem176a, Icos | Early decrease then stable |
| NK | Ccl5, Gzma, Nkg7, AW112010, Ncr1, Prf1, Klrd1, Ccl4, Lgals1, Klra4 | Transient early decrease |
| Interstitial mac | C1qa, Apoe, C1qb, C1qc, Pf4, Retnla, Cd74, H2-Aa, H2-Ab1, Selenop | Consistent decrease |
| Alveolar mac | Spp1, Chil3, Ear2, Lpl, Fabp1, Ctsd, Ear1, Plet1, Abcg1, Mrc1 | Transient early increase |
| Neutrophil | S100a9, S100a8, Retnlg, Wfdc17, Ifitm1, Il1b, BC100530, Stfa2l1, Hp | Consistent increase |
| Classical mono | Fn1, Plac8, Lyz2, S100a4, Ifitm3, F13a1, Ms4a6c, Thbs1, Ifi27l2a, Ly6c2 | Transient mid increase |
| Intermediate mono | Plac8, Ifitm3, Gngt2, S100a4, Ms4a6c, Cd74, Apoe, Ifi27l2a, H2-Ab1, Lyz2 | Transient mid increase |
| Non-classical mono | Gngt2, Pou2f2, Ace, Ifitm3, Apoe, Plac8, Stap1, Sat1, Clec4a3, Spn | Transient mid increase |
| cDC1 | Cst3, H2-Ab1, H2-Aa, Cd74, Naaa, Irf8, Plbd1, Tbc1d4, H2-DMa, Cd207 | Transient early increase |
| cDC2 | H2-Ab1, Cd74, H2-Aa, Cst3, Mgl2, Ccl17, Tnip3, Cd209a, Mt1, Tmem176a | Transient early increase |
| pDC | Bst2, Siglech, Tcf4, Irf8, Ctsl, Klk1, Cd209d, Atp1b1, Rnase6, Cox6a2 | Transient early increase |
| Ccr7 DC | Ccl5, Ccl22, Fscn1, Ccl17, Basp1, Ccr7, Tbc1d4, Il12b, Rgs1, Tmem123 | Transient early increase |
| Proliferative | Hist1h1b, Stmn1, Top2a, Mki67, Tubb5, Pclaf, Tuba1b, H2afz, Hist1h4d | Transient early increase |

| Table S2: Tissue-resident macrophage markers and population structure dynamics (Related to Figure 2) | | |
|---|---|---|
| **Interstitial Macrophages** | | |
| Antigen | H2-Aa, H2-Ab1, Cd74, Cd52, H2-DMa, H2-DMb1, Ccr2, H2-Eb1, Clec4b1 | Transient mid decrease |
| Mrc1 | F13a1, Folr2, Apoe, Selenop, Mrc1, Maf, Stab1, Gas6, Pf4, Wwp1 | No change |
| Crip1 | S100a10, Crip1, Vim, Tppp3, S100a4, S100a6, Lmna, Lgals3, Anxa2 | Transient mid increase |
| **Alveolar Macrophages** | | |
| Inflammatory | Tnf, Egr1, Cxcl2, Nfkbia, Cxcl1, Tnfaip3, Il1b, Il1a, Nfkbiz, Ier2 | Transient mid increase |
| Ccl4 | Ccl4, Ccl3, Egr2, Cxcl2, Tnf, Rasgef1b, Nfkbia, Ccrl2, Il1a, Cxcl10 | No change |
| Homeostatic | Txnip, Fau, Eef1b2, Fabp1, Cdc42ep3 | Transient mid decrease |
| Lipid | Gpnmb, Ccl6, Psap, Ctsd, Cd36, Vat1, Cstb, Cd63, Ctsk, Apoe | Transient mid decrease |
| Allergy | Ear1, Ear2, Ang4, Ang5, Cst3, Lyz2, Cyba, Fcer1g, Cd9, Ear10 | No change |
| Mt | Mt1, Mt2, Ftl1, Prdx1, Txnrd1, Hmox1, Fabp5, Srxn1, Lipa, Clec4n | No change |
| Antigen | Cd74, H2-Aa, H2-Ab1, H2-DMb1, Epcam, H2-DMa, S100a6, Cd52, Fn1 | Transient early decrease |
| Proliferative | Mki67, Stmn1, Top2a, Hist1h1b, Hmgb2, Pclaf, Cenpf, Birc5, Tubb5 | No change |

**Table S3: NMF25 signature genes** (Related to Figure 3)

| *Gene* | *Cytokine* | *TF* | *Cytosol* | *Receptor* |
|---|---|---|---|---|
| Cxcl2 | X | | | |
| Cxcl1 | X | | | |
| Tnf | X | | | |
| Ccl3 | X | | | |
| Il1b | X | | | |
| Il1a | X | | | |
| Egr1 | | X | | |
| Junb | | X | | |
| Klf6 | | X | | |
| Nfkbia | | | X | |
| Plek | | | X | |
| Serpine1 | | | X | |
| Hspa8 | | | X | |
| Tnfaip2 | | | X | |
| Tnfaip3 | | | X | |
| Ubc | | | X | |
| Nlrp3 | | | X | |
| Mcl1 | | | X | |
| Cd14 | | | | X |
| Plet1 | | | | X |
| Ccrl2 | | | | X |
| Tlr2 | | | | X |

| **Table S4: Peripheral myeloid markers and population structure dynamics** Related to Figure 4) | | |
|---|---|---|
| **Monocytes** | | |
| Classical | Fn1, F13a1, Ly6c2, Thbs1, Ccr2, Ptgs2, Wfdc17, S100a10, Vim, Cd14 | Mid increase then stable |
| Intermediate | H2-Ab1, H2-Aa, Cd74, Tmem176b, Tmem176a, H2-DMb1, Aif1, H2-DMa | No change |
| Non-Classical | Pglyrp1, Ace, Cd9, Cst3, Spn, Pou2f2, Stap1, Eno3, Cd300ld, Cd300e | Mid decrease then stable |
| M-MDSC | Wfdc21, Saa3, Lcn2, Il1r2, Lrg1, Cd33, Orm1, Cd38, Net1, Vcan | Week 15-specific |
| **Dendritic Cells** | | |
| cDC1 | Ppt1, Naaa, Cd207, Sept3, Xcr1, Itgae, Rab7b, Btla, Plbd1, Wdfy4 | Consistent decrease |
| cDC2 | Epcam, S100a6, Ccl17, S100a4, Mgl2, Fxyd5, Ywhah, Fth1, Klrd1, Ccnd2 | No change |
| IFN | Ifitm6, F13a1, Cx3cr1, Plac8, Cd7, Ms4a4c, Ifitm2, Ifitm3, Cd300a, Ms4a6c | Late increase |
| Ccr7 DC | Ccl5, Ccl22, Fscn1, Ccr7, Il12b, Fabp5, Tmem123, H2-M2, Samsn1, Cd63 | No change |
| pDC | Bst2, Siglech, Plac8, Ly6d, Ctsl, Tcf4, Atp1b1, Klk1, Cox6a2, Cd209d | Transient mid increase |
| Proliferative | Hist1h1b, Hmgb2, Stmn1, Top2a, Mki67, Pclaf, Tubb5, Hist1h4d, Tuba1b, Birc5 | No change |
| **Neutrophils** | | |
| Mature | Fos, Egr1, Zfp36, Malat1, Ier2, Lst1, Junb, Fgl2, Txnip, Jun | Mid decrease then stable |
| Transitional | Wfdc17, S100a9, S100a8 | Consistent increase |
| Immature | Camp, Ngp, Retnlg, Lcn2, Lyz2, Ifitm6, S100a6, Prok2, Wfdc21, Ly6g | Consistent increase |
| Inflammatory | G0s2, Cxcl2, Nfkbia, Ccrl2, Gadd45b, Pim1, Nlrp3, Cd14, Il1b, Tnfaip2 | Transient mid increase |
| Cxcl3 Inf | Cxcl3, Cxcl2, Nfkbia, Srgn, Cebpb, Cd14, G0s2, Lmnb1, Acod1, Il1b | Transient mid increase |
| Ccl4 Inf | Ccl4, Ccl3, Cxcl2 | No change |
| Saa3 Inf | Saa3, Igfbp6, Ifitm1, Plac8, Slpi | Late increase |
| Id2 Mature | Chil3, Id2, Klhl6, Prnp, Stfa2, Ly6e, Mt1, Ptpn22, BC100530, Rpl34 | Week 15-specific |
| Saa3 Immature | Saa3, Stfa2, BC100530, Stfa2l1, Chil3, Mmp8, Stfa3, Mt1, Id2, Ly6e | Week 15-specific |
| Sfta Immature | BC100530, Stfa2, Stfa3, Gm5483, Ngp, Stfa2l1, Ifitm6, Lcn2, Mmp8, Chil3 | Week 15-specific |

| **Table S5: Lymphocyte markers and population structure dynamics** Related to Figure 6) | | |
|---|---|---|
| **T-cells** | | |
| CD4 naive | Lef1, Satb1, S1pr1, Igfbp4, Tcf7, Crlf3, Rflnb, Npc2, Ccr7, Trib2 | No change |
| CD8 naive | Cd8b1, Cd8a, Fam241a, Ccr9, Runx3, Cd7, Dapl1, Ccr7 | Mid decrease then stable |
| CD8 memory | Ly6c2, Ctla2a, Ccl5, Nkg7, Il2rb, Ctsw, Xcl1, Lyst, Klrd1, Klrk1 | Mid increase then stable |
| CD8 effector | Ccl5, Nkg7, Ccl4, Zeb2, Cx3cr1, Fasl, Klrc2, Klrd1, Klrc1, Eomes | Late increase |
| Treg | S100a6, Ikzf2, Ifi27l2a, Ctla4, S100a4, S100a10, Tnfrsf4, Srgn, Maf, Foxp3 | Transient mid increase |
| CD4 Cd29 | Itgb1, S100a10, Crip1, S100a4, Ahnak, Tmsb4x, Ly6a, S100a13, Prr13, Vim | Late increase |
| γδT/ILC | Lmo4, Tmem176a/b, S100a4, Ramp1, Cxcr6, Fosb, Lgals3, Ccr2, Rora | No change |
| Proliferative | Hist1h1b, Hmgb2, Top2a, Mki67, Pclaf | No change |
| **NK cells** | | |
| Immunomodulatory | Ctla2a, Emb, Cd27, Ltb, Tcf7, Cd7, Shisa5, Ly6e, Ccr2, Pdcd4 | Mid decrease then stable |
| Cytotoxic | Gzma, Prf1, Gzmb, Actb, Lgals1, Cma1, Irf8, Fcer1g, Nkg7, Cd52 | Mid increase then stable |
| Ccl cytotoxic | Ccl4, Ccl3, Nfkbia, Irf8, Gadd45b, Srgn, Pim1, Btg2, Icam1, Btg1 | Transient early decrease |
| Proliferative | Lig1, Pclaf, Hells, Clspn, Dhfr, Dut, Mcm3, Gmnn, Chaf1a, Mki67 | Mid increase then stable |
| **B-cells** | | |
| Naive | H2-Aa, Ighd, H2-Ab1, Fcer2a, Mef2c, Sell, Junb | Transient early decrease |
| Ighm | Iglc1, Ighm, Vpreb3, H3f3a, Iglc2, Cd79b, Ly6d, Ptma, Cd24a, Actb | Transient early increase |
| Breg | Plac8, Ly6a, Ccnd2, Itgb1, Eef1b2, Iglc1, S100a6, Npm1, Mzb1, Marcks | Early increase then stable |
| Crip1 | Crip1, Tagln2, Vim, Klf2, Ezr, Emp3, Ahnak, S100a10, Rhob, Klf6 | Transient mid increase |
